# Network analysis of promoter interactions reveals the hierarchical differences in genome organisation between human pluripotent states

**DOI:** 10.1101/2019.12.13.875286

**Authors:** Peter Chovanec, Amanda J. Collier, Christel Krueger, Csilla Várnai, Stefan Schoenfelder, Anne Corcoran, Peter J. Rugg-Gunn

**Affiliations:** Lymphocyte Signalling Programme, Babraham Institute, Cambridge, CB22 3AT, U.K.; Nuclear Dynamics Programme, Babraham Institute, Cambridge, CB22 3AT, U.K.; Epigenetics Programme, Babraham Institute, Cambridge, CB22 3AT, U.K.; Centre for Computational Biology, University of Birmingham, B15 2TT Birmingham, U.K.; Wellcome – MRC Cambridge Stem Cell Institute, Cambridge CB2 1QR, U.K.

## Abstract

A complex and poorly understood interplay between 3D genome organisation, transcription factors and chromatin state underpins cell identity. To gain a systems-level understanding of this interplay, we generated a high-resolution atlas of annotated chromatin interactions in naïve and primed human pluripotent stem cells and developed a network-graph approach to examine the atlas at multiple spatial scales. Investigating chromatin interactions as a network uncovered highly connected hubs that changed substantially in interaction frequency and in transcriptional co-regulation between pluripotent states. Small hubs frequently merged to form larger networks in primed cells, often linked by newly-formed Polycomb-associated interactions. Importantly, we identified state-specific differences in enhancer activity and interactivity that corresponded with widespread reconfiguration of transcription factor binding and target gene expression. These findings provide multilayered insights into the gene regulatory control of human pluripotency and our systems-based network approach could be applied broadly to uncover new principles of 3D genome organisation.

## INTRODUCTION

The spatial organisation of chromosomes instructs dynamic processes during development that underpin genome regulation and transcriptional control (Bonora et al., 2014; Gorkin et al., 2014; Phillips-Cremins, 2014; Rowley and Corces, 2018; Schoenfelder and Fraser, 2019). Mammalian genomes are compartmentalised into a nested hierarchy of structural features (Bouwman and de Laat, 2015; Gibcus and Dekker, 2013). Interactions between sub-chromosomal domains form two major types of spatial compartments referred to as A and B (Lieberman-Aiden et al., 2009; Rao et al., 2014) and chromatin contacts are more frequent between regions within the same compartment type (Imakaev et al., 2012; Pombo and Dillon, 2015). Further partitioning of the genome creates megabase-scale topologically associating domains (TADs) that are largely invariant across cell types, and smaller, nested sub-TADs, chromatin loops and insulating neighbourhoods that are frequently reorganised between cellular states (Bonev et al., 2017; Dixon et al., 2012; Dowen et al., 2014; Nora et al., 2012; Pękowska et al., 2018; Phillips-Cremins et al., 2013; Rao et al., 2014; Sanyal et al., 2012; Sexton et al., 2012; Stadhouders et al., 2018).

Constrained within the larger structural features, distal regulatory elements, such as enhancers, interact with target gene promoters through DNA looping to control promoter activity. Distal elements can loop to several genes, and genes can interact with multiple distal elements, resulting in complex three-dimensional interaction networks that are often re-wired upon cell state change (Beagrie et al., 2017; Freire-Pritchett et al., 2017; Sanyal et al., 2012; Schoenfelder and Fraser, 2019). The formation of regulatory interactions is directed in part by the cell type-specific occupancy of chromatin proteins and transcription factors (Furlong and Levine, 2018). The binding of these factors is often sensitive to epigenetic marks such as DNA methylation, and thus the close interplay between epigenetics and chromatin topology is critical for the appropriate gene regulatory control of cell state transitions. Investigating how chromatin interactions track with a changing epigenetic and gene regulatory landscape is important for understanding the principles of genome organisation and transcriptional control.

One of the most striking periods of epigenome reorganisation occurs as pluripotent stem cells (PSCs) transition from a naïve state to a primed state. During this transition, DNA methylation levels rise from ∼20% to ∼70% genome-wide and there is a dramatic gain in promoter-associated H3K27me3 at several thousand genes (Ficz et al., 2013; Habibi et al., 2013; Hackett et al., 2013; Leitch et al., 2013; Marks et al., 2012; Takashima et al., 2014; Theunissen et al., 2014). These events recapitulate similar molecular transitions as embryos progress from pre- to post-implantation stages of development (Auclair et al., 2014; Borgel et al., 2010; Liu et al., 2016; Smith et al., 2012) and PSCs therefore provide a tractable cell model to investigate these processes. In mouse PSCs, global epigenetic changes occur in parallel with the re-organisation of gene regulatory interactions that take place within largely preserved structural domains (Beagan et al., 2017; Pękowska et al., 2018). For example, the acquisition of H3K27me3 at gene promoters during the transition from a naïve to a primed state is concomitant with the emergence of a long-range network of promoter-promoter interactions that connect the H3K27me3 marked regions (Joshi et al., 2015). This spatial network is thought to constrain the transcriptional activity of developmental genes (Schoenfelder et al., 2015a). In human PSCs, the mapping of a subset of DNA interactions that are cohesin-associated revealed that the positioning of long-range structural loops is similar between naïve and primed states, suggesting that their TAD and insulated neighbourhood structures are largely preserved (Ji et al., 2016). Within those domains however, individual interactions connecting active enhancers to cell type-specific genes are re-wired between naïve and primed states. This raises the possibility that nested within TADs and cohesin-mediated loops, altered interactions between genes and their regulatory elements have widespread effects on controlling transcriptional changes in pluripotent state transitions. There is a pressing need, therefore, to compare regulatory interactions between enhancers and their target genes in human PSC states globally and at high resolution, and to use these data to unravel the nested structures of genome organisation.

Chromatin conformation capture technologies such as Hi-C can reveal genome-wide spatial DNA interactions (Lieberman-Aiden et al., 2009) and can be combined with sequence enrichment to increase coverage of interactions at genomic features such as promoters (promoter-capture Hi-C; PCHi-C) (Mifsud et al., 2015; Schoenfelder et al., 2015b). The rapid progress in experimental methods requires new ways to maximise the discovery of new insights from the resulting interaction data (Dekker et al., 2013). Network approaches have been applied successfully to global Hi-C data by using network modularity to identify communities of interacting loci that represent high-level chromatin features such as TADs and sub-TADs (Chen et al., 2016; Norton et al., 2018; Yan et al., 2017). A current challenge arises because capture-based Hi-C data have regions of strong local enrichment surrounded by a relatively sparse interaction matrix, precluding conventional Hi-C heatmap-based visualisation and limiting the visualisation of these data to arc diagrams or circular plot formats. This focused viewpoint restricts data examination to individual regions of interest and can miss larger, more complex spatial networks and their dynamics. Thus, the current lack of computational approaches that can represent high-resolution, capture-based DNA interaction data at a network scale hinders progress in this area.

To understand the hierarchical differences in three-dimensional genome organisation between naïve and primed human PSCs, we used Hi-C and promoter-capture Hi-C to generate a high-resolution atlas of ∼130,000 DNA interactions in these two cell states. By developing a new computational approach to visualise the atlas at a network level, we identify striking differences in organisational features between the two cell states at multiple spatial scales. These findings uncover novel insights into the gene regulatory control of human development and pluripotency. Importantly, our experimental and computational framework establish a systems-based approach that has the potential to uncover new principles of 3D genome organisation and function in a broad range of cellular contexts.

## RESULTS

### Promoter interaction mapping in naïve and primed human PSCs

We used PCHi-C to profile the global, high-resolution interactomes of 22,101 promoters in isogenic human naïve and primed PSCs. There was a strong concordance in pairwise interaction read counts between the biological replicates of the same cell type (r^2^>0.95; Figure S1A); therefore we merged replicates for all downstream analyses. PCHi-C data normalisation and signal detection using the CHiCAGO pipeline (Cairns et al., 2016) identified 75,091 significant *cis*-interactions between baited promoters and other genomic regions in naïve PSCs, and 83,782 in primed PSCs (Figure S1B). Just under half of the interactions were common to both cell types (n=39,360). As expected, *trans*-interactions represented a small minority of promoter interactions (354 interactions). In both cell types, the majority of significant interactions were between the promoters of protein-coding genes and non-promoter genomic regions (Figure S1B). Processed datasets from this large-scale resource are available through the Open Science Framework (https://osf.io/XXXXX) and Table S1, and raw sequencing reads have been deposited to the Gene Expression Omnibus (accession GSEXXXXXX).

### Network visualisation of promoter interactomes

We next developed a computational approach called *Canvas* (Chromosome architecture network visualisation at scales) to visualise high-resolution, capture-based DNA interaction data at a network scale. Network graphs were constructed where each node of the network represents an individual *HindIII* genomic fragment (average size, 4 kb) and each edge represents a significant interaction between nodes (Figure 1A). We combined all significant interactions detected in naïve and primed PSCs to produce a single, unified network graph, which retains information about whether an interaction is shared or cell type-specific (Figure 1A). The combined network was visualised with a force directed layout (Jacomy et al., 2014) that positions highly interacting nodes closer together and pulls less interacting nodes apart (Figure 1B). This network graph approach immediately revealed structural features of the promoter interaction data. For example, several gene families such as olfactory receptor genes and keratin genes form large clusters of highly interacting genomic regions (Figure 1B).

**Figure 1:**
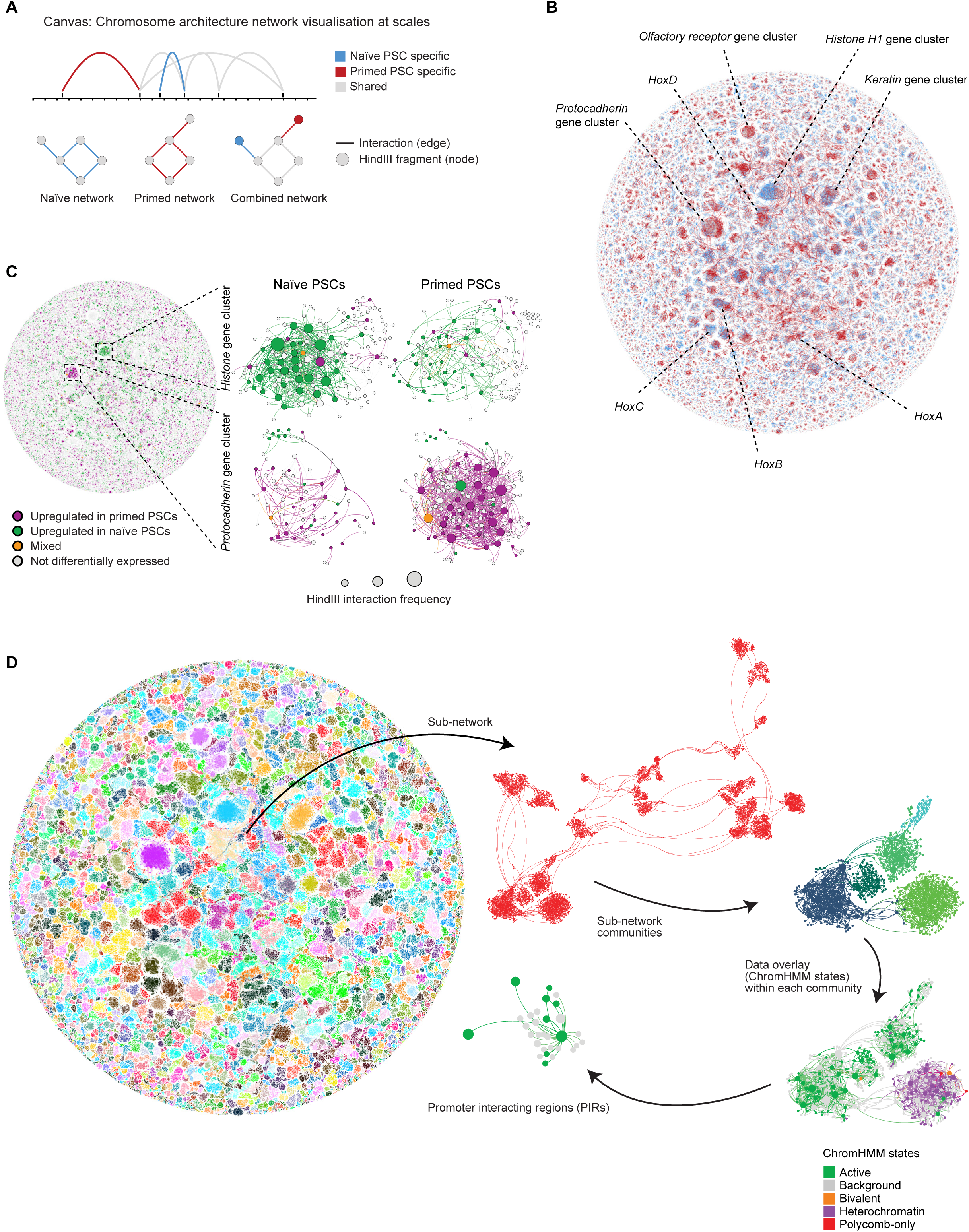
Multi-scale exploration of promoter-interaction data using force-directed network graphs. (A) Representation of PCHi-C data as arc diagrams (upper) and as corresponding network graphs (lower). Interacting *HindIII* genomic fragments are depicted as nodes that are connected by edges (significant interactions). A combined network graph is created by merging naïve and primed human PSC datasets whilst retaining cell type-specific information. Blue, naïve-specific nodes and edges; red, primed-specific nodes and edges; grey, shared nodes and edges. (B) *Canvas* produces a force-directed layout of the combined, whole-network graph. Nodes that interact more frequently are pulled closer together, and less interacting nodes are pushed further apart. (C) Differential gene expression (p-adj<0.05) categories between naïve and primed PSCs overlaid onto the combined network graph. The ‘mixed’ category refers to the small subset of nodes that contain two or more genes that differ in the direction of their transcriptional change. Expanded examples are shown for the histone H1 gene cluster (left) and the protocadherin gene cluster (right). The size of each node corresponds to the number of interactions (degree) of the node. (D) *Canvas* enables a multi-scale investigation of genome organisation. See also Figures S1 and S2.

Annotating naïve-specific and primed-specific interactions onto the combined network uncovered large clusters with high and uniform cell type-specific interactivity. Two of the most prominent examples of this are the histone H1 genes in naïve PSCs and the protocadherin genes in primed PSCs (Figures 1B and S2A). The histone H1 cluster contained 198 nodes connected by 984 edges, and the protocadherin cluster contained 165 nodes and 1188 edges. Overlaying transcriptional information onto each node showed that the higher promoter interactivity within the histone H1 and in the protocadherin clusters is associated with increased expression of nearly all genes within each cluster, implying a coordinated transcriptional response (Figures 1C and S2B). Interestingly, not all large gene clusters that changed interaction frequency between cell types showed differential expression, as exemplified for the keratin and olfactory receptor gene regions (Figures 1B, S2A and S2B). Taken together, the combination of high-resolution DNA interactivity and gene expression information on a whole network graph allows a systems-level visualisation of genome organisation features that might otherwise have been missed by conventional methods.

### Multiscale exploration of promoter interaction networks

Network reconstruction of the promoter-interaction data allows the interrogation of genome organisation at multiple levels. Viewed at the lowest magnification, the network graph consists of >3000 individual sub-networks of varying size (Figure 1D). Each sub-network is further divided into communities that have high internal interactivity, using algorithms developed for the detection of community structure (Blondel et al., 2008). Furthermore, overlaying histone modification information onto the network can additionally reveal common chromatin states within groups of interacting genes (Figure 1D). The highest visualisation level identifies detailed structure including potential *cis*-regulatory regions of individual baited gene promoters (Figure 1D).

### Infrequent spatial compartment switching between naïve and primed human PSCs

We first used the network graph to investigate the high-level topological features of naïve and primed PSCs. We generated matched Hi-C datasets to define the locations of TADs and assigned each node in the network to a specific TAD. Projecting the TADs on to the network graph showed that the majority (70%) of TADs are shared between the two cell types (Figure 2A). Although the positioning of TADs was similar, the insulation score of TAD boundaries was slightly higher in primed compared to naïve PSCs (Figure 2B), which suggests there are differences in TAD boundary strength between the two cell types. Interestingly, we noticed that the positioning of the TADs showed a strong, significant overlap with the individual communities that were identified within the promoter-interaction data (Figures 2C and 2D). This finding demonstrates that *Canvas* is capable of defining structural features of genome organisation.

**Figure 2:**
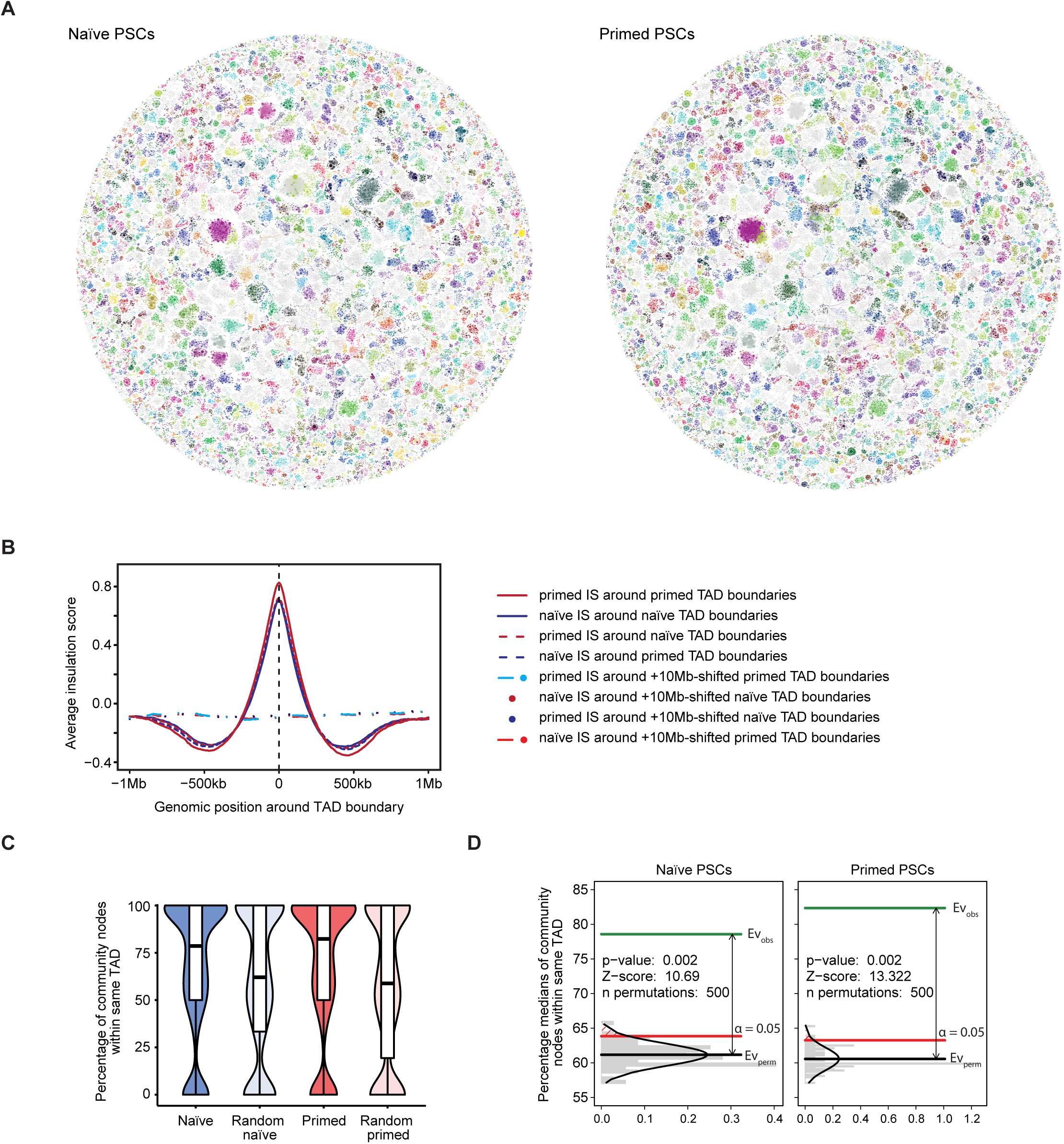
Identified communities recapitulate TADs and are largely shared between naïve and primed PSCs. (A) TADs were defined using matched Hi-C data and their locations are projected onto the PCHi-C network graph. Each TAD is represented by a unique colour and TADs shared between naïve and primed PSCs have the same colour. Nodes that fall outside of a defined TAD are coloured in grey. (B) Insulation score (IS) profiles around TAD boundaries in naïve and primed PSCs. The dotted lines show the IS around boundaries that have been computationally shifted by +1 Mb. (C) Plot (left) shows the percentage of nodes within each community that are contained within the same TAD. For both cell types, the percentage was significantly higher than compared to a set of randomly shuffled TAD coordinates. (D) Statistical test for (C). Bar chart shows the frequency of median percentages obtained after 500 random permutations of TAD coordinates (grey) and the comparison between this control set (Ev_perm_) and the observed values (Ev_obs_) using a permutation test within the regioneR package (Gel et al., 2016).

We next examined spatial compartmentalisation in naïve and primed PSCs. Annotating each node with eigenvalue scores and chromatin mark profiles segregated the genomic regions into ‘A’ (euchromatin) and ‘B’ (heterochromatin) compartments (Figure 3A). Consistent with previous work that compared compartments between other cell types, the majority of interacting regions are in the same compartment in naïve and primed PSCs (88%; n=57329). Of the interacting regions that do change compartments between the two cell types (12%; n=7956), there are roughly equal numbers that transition in each direction. Figure 3B shows examples of gene promoters such as *TET2* and *JAK2* and their interacting regions that switch compartments and are transcriptionally altered. However, for the majority of regions, compartment switching and transcriptional changes were not correlated, as exemplified for *HNRNPK* and *KIF27*, which are upregulated in primed PSCs despite switching from an ‘A’ compartment in naïve PSCs to a ‘B’ compartment in primed PSCs (Figure 3B). These results indicate that changes in transcription and compartmentalisation can be uncoupled, in line with previous studies (Stadhouders et al., 2018). Although the majority of compartments were common to both cell types, the compartmentalisation strength of naïve PSCs was substantially lower than for primed PSCs (Figure 3C). This finding implies that naïve PSCs show a higher degree of variation in genome conformation or that the strength of compartmental attraction differs between the two pluripotent states (Nuebler et al., 2018). Taken together, network scale visualisation of promoter interactomes and chromatin state annotation can provide an intuitive method for data exploration including at the level of higher-order chromatin organisation.

**Figure 3:**
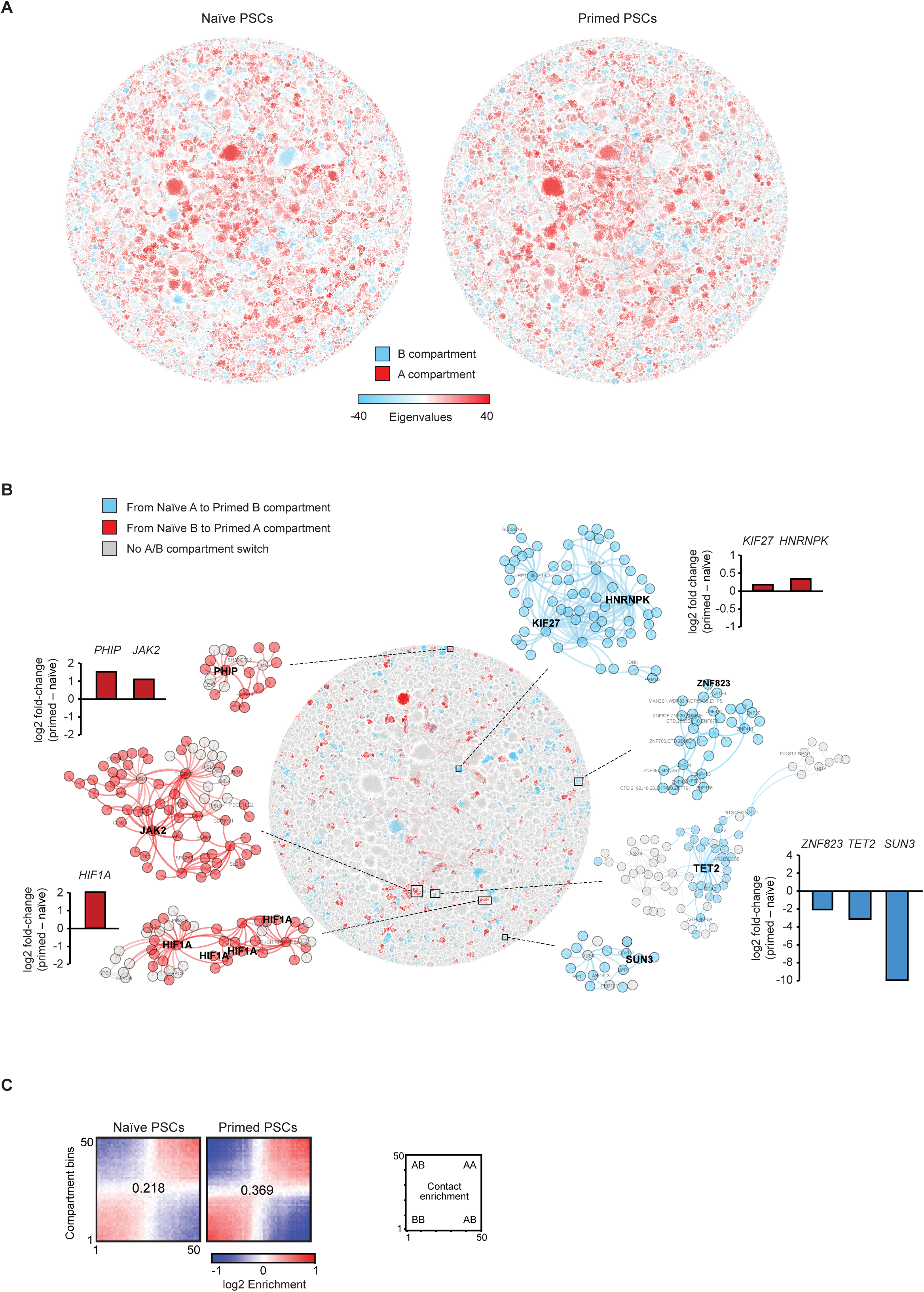
Compartmentalisation strength, but not A/B identity, differs between naïve and primed PSCs. (A) Overlay of PC1 Eigenvalues onto the PCHi-C network graphs annotates each node to ‘A’ or ‘B’ compartments. (B) Examples of nodes that undergo A/B compartment switching between naïve and primed states. The nodes are coloured according to the direction of compartmentalisation switch. Column charts show the fold-change in transcription levels between naïve and primed PSCs for the examples shown. (C) Heatmap contact matrix shows the compartmentalisation strength in naïve and primed PSCs. Genomic regions (250 kb) were allocated into 50 bins according to their principal component values ordered from the most ‘B-like’ (#1) to the most ‘A-like’ (#50). Using the Hi-C data, the log2 contact enrichment between each of the bins are shown as a heatmap matrix. Bins within the same compartment interact more frequently, as expected. However, the log2 contact enrichment score between bins of the same compartment is higher in primed compared to naïve PSCs, and the overall compartmentalisation strength is also higher (values are shown in the centre of each matrix).

### Long-range promoter interactions distinguish the two pluripotent states

We next examined promoter interaction changes between pluripotent states by comparing the number of nodes and the overall number of interactions within each individual sub-network. Using this approach, the protocadherin and histone H1 clusters stood out as prominent outliers (Figure 4A). More generally, sub-networks in primed PSCs tended to be larger and contain a greater number of interactions and with more nodes (Figure 4A). These findings suggest that smaller communities in naïve PSCs come together to form larger sub-networks in primed PSCs.

**Figure 4:**
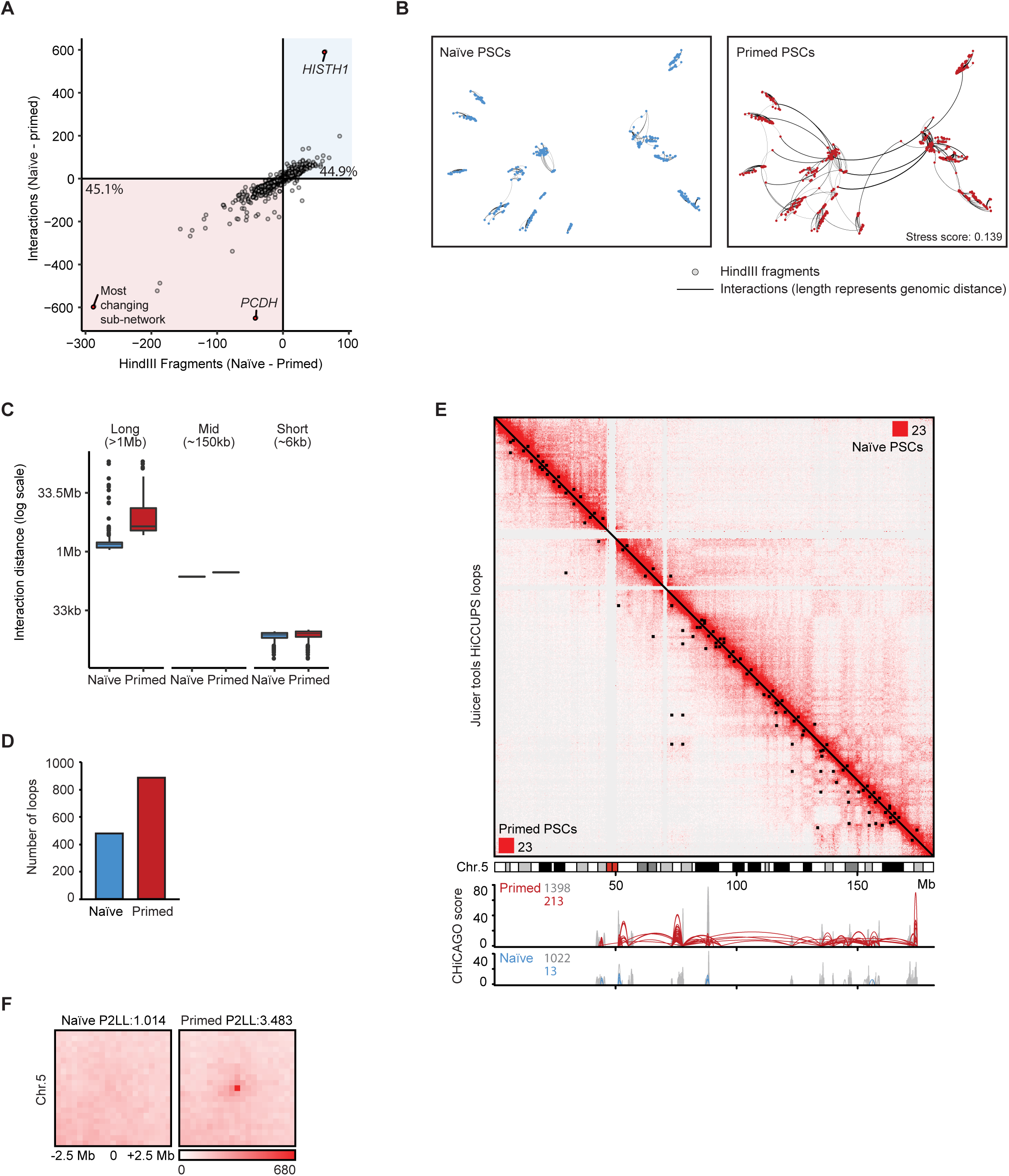
Long-range promoter interactions in primed PSCs drive genome conformation changes between pluripotent states. (A) Plot shows the number of interactions (edges) and the number of interacting *HindIII* fragments (nodes) for each sub-network in naïve and primed PSCs. Each small circle represents a different sub-network. The lower-left quadrant contains larger sub-networks in primed PSCs, and the upper-right quadrant contains larger sub-networks in naïve PSCs. The protocadherin (*PCDH*), histone H1 (*HISTH1*) and ‘most changing’ (containing diverse genes) sub-networks are highlighted in red. (B) Multidimensional scaling representation (MDS) of the ‘most changing’ sub-network plotted using the linear genomic distance between nodes as edge weights. The measured stress score of 0.139 indicates there is a reasonable fit between the linear genomic distances and the spacing of the nodes as determined by MDS (Kruskal and Wish, 1978). (C) Plot shows the distribution of linear genomic distances between interacting nodes in naïve and primed PSCs, binned into long-, mid-, and short-range distances. (D) Chart shows the total number of chromatin loops identified on chromosome 5 in naïve and primed PSCs. (E) Hi-C interaction matrix of chromosome 5 at a resolution of 250 kb with Knight-Ruiz (KR) normalisation; upper right, naïve PSCs; lower left, primed PSCs. Areas of contact enrichment were defined separately for naïve and primed PSCs using HiCCUPS and each cell type-specific set of chromatin interactions are highlighted as a black square on their respective heatmap. The two corner numbers indicates the maximum intensity values for the matrix. The tracks below the Hi-C heatmap show the PCHi-C interactions and ChiCAGO scores over the same region. (F) Heatmap shows the aggregate peak analysis (250 kb resolution) of primed-specific chromatin interactions on chromosome 5 for naïve and primed PSCs. Chromatin interactions >5 Mb from the diagonal were used for the analysis (n=27 loops out of a total of 76). The ‘peak to lower left’ (P2LL) score denotes the enrichment of the central pixel over the pixels in the lower left quadrant. See also Figures S3 and S4.

To begin to understand the increased size of sub-networks in primed PSCs, we examined the sub-network that showed the largest change in the number of nodes and edges when comparing between pluripotent states. In naïve PSCs, this sub-network was composed mainly of individual, separated communities that are distributed across chromosome 5 (Figure 4B). However, in primed PSCs, these communities were connected by >200 long-range interactions (defined as >1 Mb in linear distance) (Figure 4B). The acquisition of long-range interactions to create large sub-networks in primed PSCs was a common feature, as exemplified for regions on other chromosomes including the *HOXA*, *HOXD* and *NKX* loci (Figures S3A and S3B). In contrast, the only clear instance of a sub-network with higher numbers of promoter interactions and nodes in naïve PSCs was for the histone H1 locus (Figures S3A and S3B). Analysing all promoter interactions revealed a substantial increase in their number and linear distance in primed compared to naïve PSCs (Figure 4C) and this difference was independent of the applied CHiCAGO threshold (Figure S3C).

We next confirmed the difference in the number of long-range promoter interactions between pluripotent states using an alternative approach that is independent of promoter capture. We analysed our Hi-C data using HiCCUPS (Rao et al., 2014), which is an algorithm that calls interaction ‘peaks’ when a pair of loci show elevated contact frequency relative to the local background. This approach revealed that the number of long-range chromatin interactions (>1Mb) was substantially higher in primed (n=889) compared to naïve PSCs (n=480) (Figure 4D). This striking difference is exemplified for several chromosomes by overlaying the identified peaks onto Hi-C contact matrices (Figures 4E and S4). Importantly, the interaction peaks identified by HiCCUPS in primed PSCs matched the positions of long-range promoter interactions detected by PCHi-C (Figure 4E), thereby validating the presence of the promoter-capture interactions. To visualise the contact enrichment at the identified peaks more closely, we applied Aggregate Peak Analysis (Rao et al., 2014) to the long-range interaction loci and found there was a strong enrichment at these aggregated sites over local background levels in primed PSCs but not in naïve PSCs (Figure 4F). This result confirms the observed differences in long-range contact frequency between pluripotent cell types. Taken together, long-range promoter interactions are a dominant feature in primed PSCs that connect individual communities into larger sub-networks.

### Acquisition of Polycomb-associated interaction networks in primed human PSCs

To begin to characterise the properties of the long-range promoter interactions, we used published and newly generated histone ChIP-seq data to assign ChromHMM-defined (Ernst and Kellis, 2012) chromatin states to individual *HindIII* fragments (Figures S5A and S5B). Interestingly, the chromatin state of regions that are brought together by long-range interactions differed between the two cell types. The longest interactions in naïve PSCs were marked by active chromatin states at the promoter and at the interacting region (Figure 5A, left). These interactions and their chromatin state were also present in primed PSCs (Figure 5A, left). In contrast, the longest interactions in primed PSCs were dominated by bivalent chromatin states (dual H3K27me3 and H3K4me1/3 methylation) at both ends of the interaction (Figure 5A, right). This set of regions has reduced bivalent marks in naïve PSCs and very few of these interactions were detected in naïve PSCs, suggesting they are formed de novo upon the transition from naïve to primed pluripotency.

**Figure 5:**
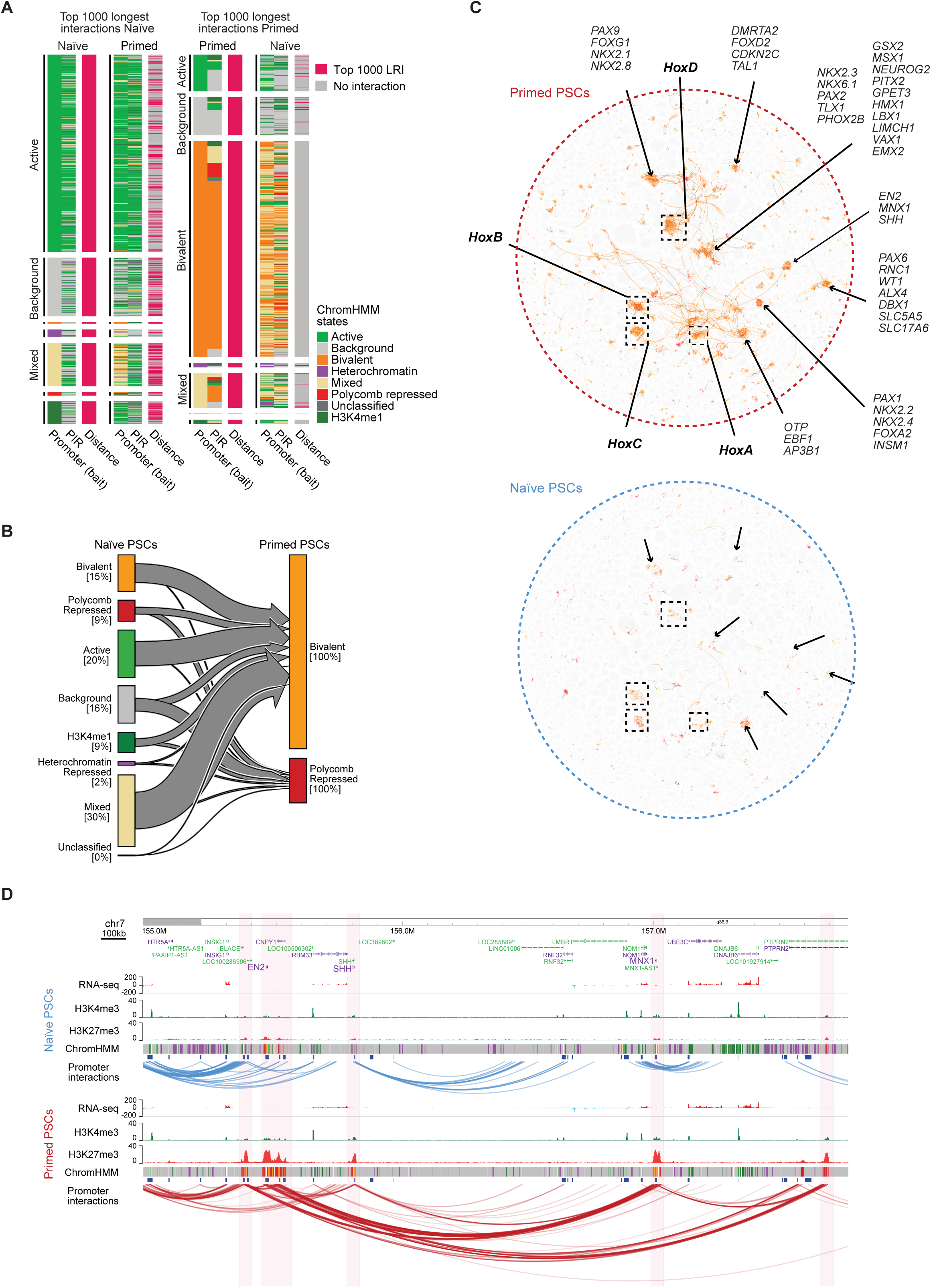
Long-range promoter interactions are associated with bivalent chromatin in primed human PSCs. (A) Heatmaps show the chromatin states of the promoters and promoter-interacting regions (PIRs) that are connected by the longest interactions in naïve (left) and primed (right) PSCs. Each heatmap is ordered and divided by the chromatin state of the promoter *HindIII* fragment. (B) Plot shows the chromatin states of *HindIII* fragments in naïve PSCs that transition into the bivalent or Polycomb-associated states in primed PSCs. The percentages shown are within each state. (C) Visualising all bivalent and Polycomb-associated interactions on the network graph highlights several interaction clusters particularly in primed PSCs (upper) that contain many developmental genes. The clusters, together with the interactions, are largely absent in naïve PSCs (lower). (D) Genome browser view in naïve (upper) and primed (lower) PSCs of a ∼2.5 Mb region that contains several developmental genes including *EN2*, *SHH* and *MNX1*. Significant promoter-interactions are shown as blue and red arcs. ChIP-seq (H3K4me3 and H3K27me3) and RNA-seq tracks are shown. Chromatin states for each genomic region were defined by ChromHMM (Ernst and Kellis, 2012) using ChIP-seq data (active chromatin, light green; H3K4me1-only chromatin, dark green; bivalent chromatin, orange; Polycomb-associated, red; heterochromatin, purple; background, grey). See also Figure S5.

To investigate the establishment of H3K27me3-associated interaction networks in primed PSCs, we examined the chromatin state of these same regions in naïve PSCs. Approximately one-quarter of the regions were already marked by H3K27me3 in naïve PSCs (Figure 5B). More commonly, the regions were classified as active or mixed state chromatin in naïve PSCs, and therefore acquired H3K27me3 during the transition to primed PSCs (Figure 5B). A closer look at the individual *HindIII* fragments that were classified as mixed chromatin state in naïve PSCs revealed that these regions contained patches of active and repressive chromatin states (Figure S5C), commonly with H3K4me1/3-only peaks residing within larger blocks of H3K27me3. This implies that the H3K4me1/3-marked sites are protected from H3K27me3 in naïve PSCs, but that H3K27me3 spreads throughout the region in primed PSCs. Genes within these regions were associated with developmental processes, with examples including *DLX*, *GATA* and *HOX* factors.

Using *Canvas* to visualise all H3K27me3-associated interactions further highlighted the differences between cell types. This category of interactions formed numerous highly interacting clusters in primed but not naïve PSCs (Figure 5C). Individual clusters were connected through long-range interactions and indeed nearly all (98%) of the long-range *cis*-interactions within the data set were associated with bivalently-marked promoters (Figure S5D).

Protein-coding genes belonging to the major developmental gene families were spatially organised within the H3K27me3-associated interaction network in primed PSCs (n=696; Figure 5C). This gene set was strongly enriched for transcriptional regulators and homeobox-containing factors (Table S2). For example, a region on chromosome 7 that includes *SHH*, *EN2* and *MNX1* formed a highly interacting cluster in primed PSCs through the presence of long-range interactions that align closely to H3K27me3 peaks (Figure 5D). Other examples include the *HOX* gene loci, where we detected long-range *cis*-interactions and also *trans*-interactions with regions on other chromosomes, including the *HOX* clusters themselves (Figure S6A). The lower levels of H3K27me3 at gene promoters in naïve PSCs corresponded to the absence of interactions within the *HOX* loci, while interactions outside of the regions were similar in both cell types (Figures S6B and S6C). Taken together, these results show that the majority of long-range interactions connect regions that gain H3K27me3 during the naïve to primed conversion, thereby creating large spatial networks in primed PSCs.

### Pluripotent state-specific enhancer activity and interactivity

To investigate changes in gene regulatory control between pluripotent states, we annotated enhancers in each cell type and then used PCHi-C data to identify the target gene promoters for those enhancers. We defined super-enhancers (SEs) by running H3K27ac ChIP-seq data through the ranking of super-enhancer (ROSE) pipeline (Lovén et al., 2013; Whyte et al., 2013). This approach identified 182 naïve-specific SEs and 62 primed-specific SEs (Figure 6A). We also curated ∼600-700 SEs that are shared between both cell types (Figure 6A). Integrating the enhancer annotations with the PCHi-C data identified the gene promoters that interact with each SE. Interestingly, the majority of SE-target genes (85%, n=951) were cell type-specific and only 15% of genes were contacted by SEs in both naïve and primed PSCs (Figure 6B). Genes that interacted with SEs only in naïve PSCs included members of signalling pathways such as *IL6* and *GDF3* and chromatin regulators such as *TET1* and *REST* (Figure 6B). Similarly, genes in contact with a SE only in primed PSCs included transcription factors such as *KLF7*, *TCF4* and *ZIC2* (Figure 6B). Relatively few genes (n=176) interacted with a SE in both cell types and this set of genes included *ACTB* and *LIN28B* (Figure 6B). In line with the proposed capability of SEs to promote the expression of genes that are important for cell identity (Hnisz et al., 2013; Whyte et al., 2013), we observed a higher transcriptional output of SE-target genes in a cell type-specific manner and a comparable, high transcriptional output for genes contacted by SEs in both cell types (Figure 6C). The large number of cell type-dependent gene promoters that contact SEs underscores the substantial changes in SE activity and interactivity that occur between the two pluripotent states.

**Figure 6.**
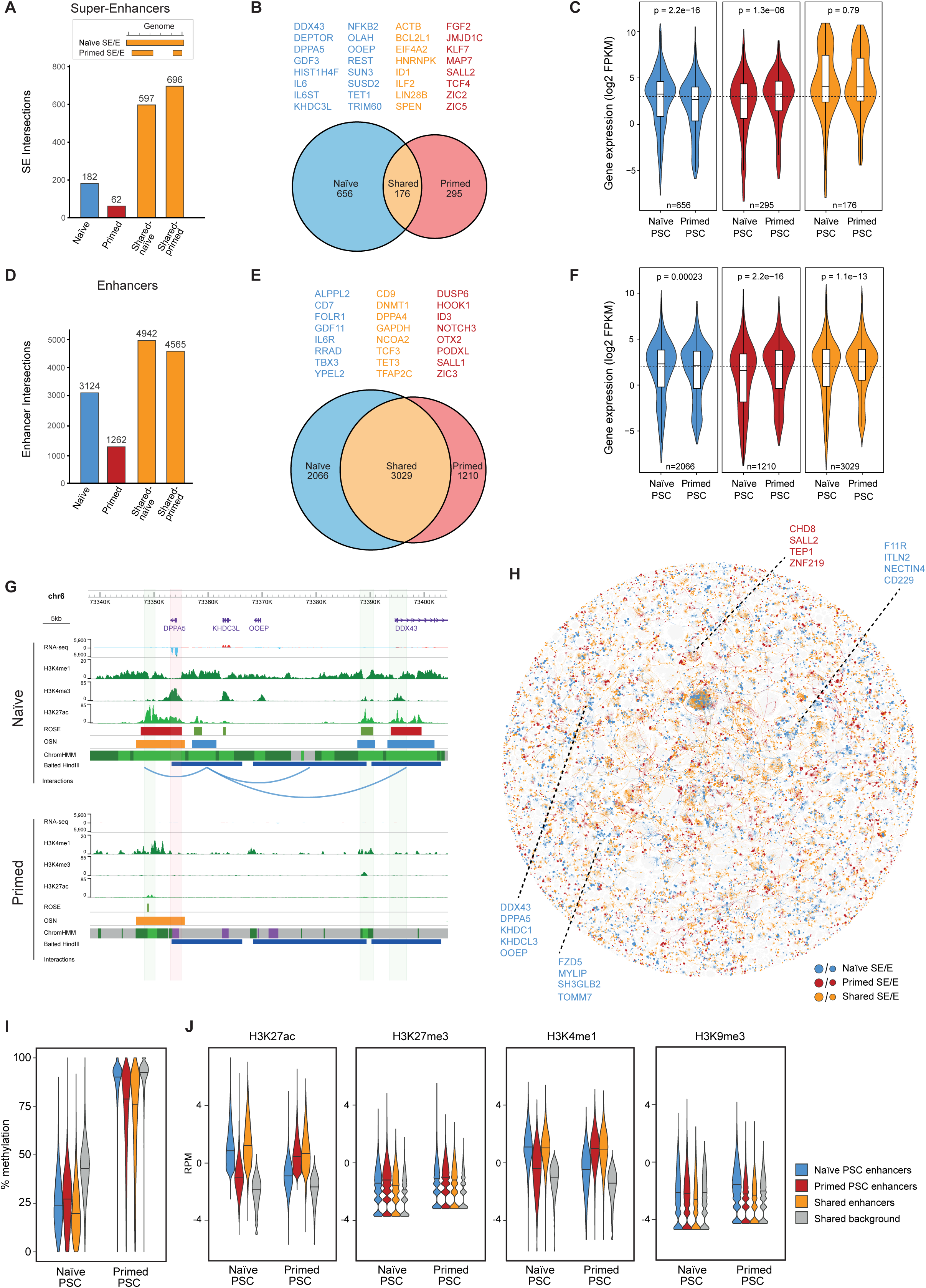
Dynamics of enhancer activity and interactivity between human pluripotent states. (A) Plot showing the number of ROSE-called SEs in naïve and primed PSCs. As illustrated in the diagram, two values are given for shared SEs because a SE in one cell type may overlap with two individually-called SEs in the other cell type. (B) Diagram showing the number of genes that are contacted by SEs in the two pluripotent cell types. Shared genes (orange) are genes that are contacted by SE elements in both naïve and primed PSCs. Naïve-specific genes (blue) and primed-specific genes (red) are contacted by SEs in either naïve or primed PSCs, respectively. (C) Plots showing the log2 FPKM expression of genes that interact with SEs in each cell type (naïve, n=648; primed, n=286; shared, n=174). P-values are derived from a Mann Whitney U test. (D) Plot showing the distribution of ROSE-called enhancers in naïve and primed PSCs. (E) Diagram showing the number of genes that are contacted by enhancers in the two pluripotent cell types. Genes that are also in contact with a SE have been removed from this list of enhancer-interacting genes. (F) Plots showing the log2 FPKM expression of genes that interact with enhancer elements in each cell type (naïve, n=2077; primed, n=1210; shared, n=3158). P-values are derived from a Mann Whitney U test. (G) Genome browser view of the *DPPA5* promoter interactomes in naïve (upper) and primed (lower) PSCs. Significant interactions are shown as blue arcs that connect the baited *HindIII* fragment containing the *DPPA5* promoter (shaded in red) with promoter-interacting regions (shaded in green). ChIP-seq (H3K4me1, H3K4me3 and H3K27ac) and RNA-seq tracks are shown. Chromatin states include active chromatin, light green; H3K4me1-only chromatin, dark green; bivalent chromatin, purple; background, grey. ROSE tracks show the location of enhancers (green) and super-enhancers (red), and OSN tracks show the position of shared (orange) and naïve-specific (blue) regions of OSN occupancy. (H) Network graph showing the locations and cell-type-origin of enhancer and SE elements. Colours depict naïve-specific (blue), primed-specific (red) and shared (orange) enhancer and SE elements. Node size represents SE (large nodes) and enhancers (small nodes). Lines represent interactions and are coloured according to the colour of the node of origin. (I-J) Plots show the (I) percent DNA methylation or (J) histone modification levels in naïve and primed PSCs at shared (n=18,735) and cell type-specific active enhancers (naïve, n=26,955; primed, n=34,805). Regions that are in the background chromatin state in both cell types are shown to indicate genome-wide levels (n=467,772). See also Figure S6.

We next examined interactions between promoters and enhancers other than SEs (Figure 6D). A high proportion (48%) of gene promoters were in contact with enhancers in both cell types (Figure 6E), which contrasts with the much lower proportion for SEs (15%; Figure 6B). Several core pluripotency-associated genes interact with enhancers in both naïve and primed PSCs including *DPPA4*, *TCF3* and *TFAP2C* – all of which are highly expressed in both cell types. More generally, differences in enhancer interactivity were concordant with transcriptional changes of their target genes in naïve and primed PSCs (Figure 6F). Genes that interact with enhancers only in naïve PSCs include *RRAD* and *YPEL2*, and in primed PSCs this gene set includes *DUSP6* and *OTX2*. Given the differences in the transcriptional activity and regulatory control of the identified genes, this integrated dataset uncovers factors that could have important cell type-specific functions.

The resultant data set uncovered changes in promoter – enhancer interactions that occur between naïve and primed PSCs, thereby revealing new insights into gene regulatory control of human pluripotency. For example, *DPPA5* is highly transcribed in naïve PSCs and the promoter interacts with three SEs that are marked by high levels of H3K27ac and H3K4me1 (Figure 6G). In contrast, there are no SEs within this region in primed PSCs, no detectable *DPPA5* promoter interactions and *DPPA5* is not transcribed (Figure 6G). A second example shown is *TBX3*, which is more highly transcribed in naïve compared to primed PSCs, and this corresponds to the presence of *TBX3* promoter interactions with enhancers only in naïve PSCs (Figure S6D).

We next took advantage of *Canvas* to examine promoter – enhancer communication on a network scale (Figure 6H). The global visualisation of promoter interactions and enhancer annotations revealed clusters of co-regulated genes with many hubs containing both SEs and enhancers (Figure 6H). Collectively, these integrated data sets, therefore, provide an important resource to investigate the changes in the activity and target gene interactivity of putative regulatory regions in human pluripotent states.

To better understand the changes in enhancer activity between human pluripotent states, we next examined how naïve-specific active enhancers are decommissioned in primed PSCs. By far the most common change at these sites was an increase in DNA methylation levels from an average of ∼25% in naïve PSCs to ∼90% in primed PSCs (Figure 6I). The gain in DNA methylation at these regions was very similar to the increase that occurs genome-wide, as indicated by the ‘background’ category and reported previously (Figure 6I; (Pastor et al., 2016; Takashima et al., 2014)). In naïve PSCs, all enhancer categories were, overall, less methylated compared to background (>10% difference in median DNA methylation levels). In primed PSCs, however, only primed-specific and shared enhancer categories showed lower DNA methylation than background (by >10%), suggesting that these regions are protected to some extent from global events (Figure 6I). Examination of histone ChIP-Seq data confirmed the expected reduction in H3K27ac and H3K4me1 levels at naïve-specific active enhancers when comparing between naïve and primed PSCs (Figure 6J, log2 fold-change of median >1). The majority of naïve-specific active enhancers did not gain H3K9me3 or H3K27me3 in primed PSCs (log2 fold-change of median <1), thereby demonstrating that the acquisition of repressive histone modifications is not a common mode of enhancer decommissioning in primed PSCs (Figure 6J). However, we identified a set of ∼100 enhancers that acquired H3K27me3 in primed PSCs and these regions were associated with developmental genes including *GATA3*, *TFAP2A* and *NEUROG1* (Figure S6E). In keeping with the known antagonism between H3K27me3 and DNA methylation processes, most of the enhancers that gained H3K27me3 remained DNA hypomethylated in primed PSCs (Figure S6F). Thus, the majority of naïve-specific active enhancers are decommissioned by acquiring DNA methylation, however a small subset of these enhancers adopt a Polycomb-associated chromatin state in primed PSCs. Taken together, this integrated data resource provides a large collection of putative regulatory sequences and their target promoters in naïve and primed PSCs, underpinned by network interactions, thereby revealing the dynamic changes in gene regulatory control between human pluripotent states.

### Widespread reorganisation in OCT4, SOX2 and NANOG occupancy between human pluripotent states

To begin to understand how the dynamic regulation of enhancers is controlled between human pluripotent states, we investigated the association between transcription factor binding and enhancer activation. After integrating new and previously generated ChIP-seq data, we observed that the shared pluripotency factors OCT4, SOX2 and NANOG (OSN) showed clear cell type-specific binding at the identified *DPPA5* promoter interacting regions, whereby OSN was bound at the interacting SEs only in naïve PSCs (Figures 6G). To study this association at a genome-wide level, we first asked whether OSN bound sites overlap with particular chromatin state categories. The results showed that OSN occupancy is strongly associated with active enhancers in both cell types (Figure 7A). Interestingly, essentially all OSN sites in both cell types contained enhancer chromatin signatures including H3K27ac and H3K4me1 signals and open ATAC-seq regions (Figure S7). Similarly, nearly all active enhancers showed OSN signal (Figures 7B and 7C). These results demonstrate a remarkable overlap between OSN occupancy and active enhancers in naïve and primed PSCs.

**Figure 7.**
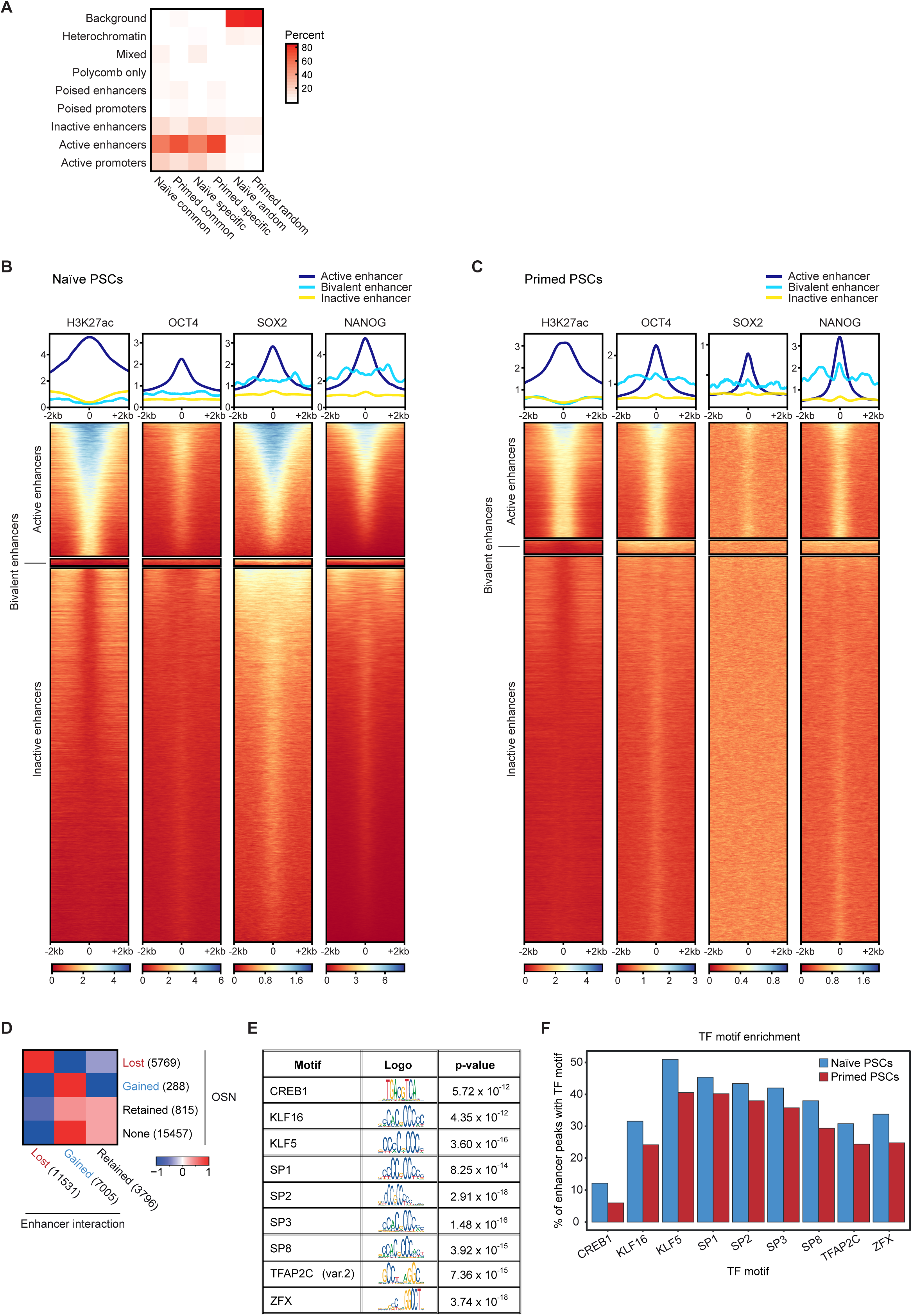
Widespread reorganisation of OCT4, SOX2 and NANOG binding at enhancers occurs between human pluripotent states. (A) Heatmap shows the percentage of OSN sites that fall within each of the ChromHMM-defined chromatin states. Columns 1 and 2 indicate OSN sites that are common to both naïve and primed PSCs; columns 3 and 4 are OSN sites that are specific to either naïve or primed PSCs; columns 5 and 6 represent random regions that do not contain OSN sites. (B–C) ChIP-seq data for H3K27ac, OCT4, SOX2 and NANOG in (B) naïve and (C) primed PSCs. Metaplots (upper) and heatmaps (lower) show normalised ChIP-seq read counts within a 4 kb peak-centered window. Regions were subsetted into active enhancers (naïve, n=44,144; primed, 51,803), bivalent enhancers (naïve, n=2,357; primed, n=6,181) and inactive enhancers (naïve, n=123,416; primed, n=173,067) based on ChromHMM-defined chromatin states, and ranked by H3K27ac signal. (D) Heatmap shows the log2 odds ratio for the associated changes in OSN occupancy and promoter-enhancer interactions in primed compared to naïve PSCs. (E) Table shows the highest-ranking (by adjusted p-value, Fisher’s exact test) transcription factor motifs that are enriched at OSN sites in naïve compared to primed PSCs. Four motifs associated with transcription factors that are not expressed in naïve PSCs (log2 RPKM < 0) were removed from the list: KLF1, NR2F1, ZNF354C and VDR. (F) Bar chart shows the percentage of OSN-bound enhancers that contain each of the identified transcription factor motifs in naïve and primed PSCs. See also Figure S7.

Despite the strong associations between OSN binding and active enhancers in both cell types, only a small proportion (<20%) of active enhancers are shared between naïve and primed PSCs (Table S3). This observation suggests there is widespread remodelling of OSN occupancy and enhancer activity between naïve and primed PSCs. Furthermore, the loss of OSN occupancy during the transition from a naïve to a primed state was strongly associated with the loss of interactivity at those regions, and vice versa for sites that gain OSN binding (Figure 7D). Enhancers that either retained OSN binding or were not bound by OSN typically retained or gained interactions (Figure 7D), which is probably due to the higher number of OSN-bound enhancers in naïve PSCs (Figure S7).

Given that OSN bind different active enhancers in each cell type, OSN is likely to be recruited by cell type-specific factors. As a first look at this, we identified transcription factor motifs that were significantly enriched at OSN sites in one cell type compared to the other. Few primed-specific candidate factors were identified, however our analysis uncovered several transcription factors that were expressed in naïve PSCs and whose binding motif occurred more frequently at OSN sites in naïve compared to primed PSCs (Figure 7E). These factors included KLF5, KLF16, SP transcription factors, TFAP2C and ZFX (Figures 7E and 7F). Thus, combinations of these transcription factors could help to recruit the shared pluripotency factors OSN to active enhancers in naïve PSCs. Taken together, these findings uncover the substantial and global reorganization in enhancer activity, interactivity and OSN-binding that occurs during the transition between human pluripotent states.

## DISCUSSION

We generated high-resolution profiles of chromatin interactions and enhancer states in naive and primed PSCs and interrogated the data using a newly developed network-scale computational approach. This viewpoint uncovered widespread rewiring particularly of large interaction sub-networks and also of promoter – enhancer contacts that change between pluripotent cell types. These findings together with the annotated chromatin interaction maps advance our systems-level understanding of the molecular control of gene regulation in pluripotency and in the earliest stages of human development.

The high-level structural organisation of TADs and spatial compartments was similar between pluripotent states, which is in agreement with recent previous reports in these and other cell types (Battle et al., 2019; Dixon et al., 2015; Ji et al., 2016). Within those topological features, however, interaction sub-networks that are formed of large, highly connected hubs changed substantially in their interaction frequency and, for a subset, also in their transcriptional activity between pluripotent states. A prominent example of this was for a region containing multiple histone H1 genes, which unexpectedly had higher promoter interactivity and transcriptional output in naïve PSCs compared to primed PSCs. This cluster included sets of histone H1 genes that are transcribed with cell cycle-dependent and independent control, which suggests that the difference in the chromatin organisation of this region between pluripotent states is likely to be driven by additional factors that act outside of the cell cycle. Given that the expression and regulation of individual H1 isoforms varies substantially between cell types and these properties are associated closely with pluripotency, differentiation and development (Pan and Fan, 2016; Turinetto and Giachino, 2015), a focused examination of histone H1 function in these cell types would be an interesting future direction of research. More commonly though, we discovered that sub-networks tended to be larger and to contain a greater number of interactions in primed compared to naïve PSCs. Examples of this included the protocadherin gene cluster, which encodes cell adhesion molecules that are predominantly expressed in the neural lineage (Chen and Maniatis, 2013). In neural cells, active protocadherin gene promoters and enhancers are brought together by CTCF and Cohesin-mediated DNA loops to form a ‘transcriptional hub’ (Guo et al., 2012). Our results show that this hub begins to pre-form during early development, potentially priming this region for co-ordinated activation upon neural development. More generally, cell-specific changes in sub-network organisation may provide opportunities for transcriptional co-regulation and resilience to perturbation. Our study has identified a large cohort of networks that can now be systematically targeted to test these predictions.

The aggregation of interaction hubs into larger networks in primed PSCs was frequently associated with the acquisition of long-range interactions that bridged Polycomb-occupied regions. These events created spatial networks connecting >600 cis-regulatory elements that control the transcription of developmental regulators. Similar, smaller-scale, Polycomb-mediated networks have been described in serum-grown mouse PSCs (Denholtz et al., 2013; Joshi et al., 2015; Schoenfelder et al., 2015a) and the prevalence of Polycomb-associated long-range interactions is strongly reduced after mouse PSCs are transitioned to a naïve state (Joshi et al., 2015). These changes have been attributed in mouse PSCs to the global redistribution of DNA methylation and H3K27me3 that occurs between primed and naïve pluripotent states (Joshi et al., 2015; Marks et al., 2012; van Mierlo et al., 2019). Interestingly, preventing the redistribution of epigenetic marks during the transition to a naïve state is sufficient to block changes in chromatin compaction at several exemplar regions, thereby directly linking epigenome remodelling with aspects of genome organisation (McLaughlin et al.). The transcriptome and cell state of mouse and human naïve PSCs are largely unaffected by experimentally disrupting Polycomb levels (Galonska et al., 2015; Moody et al., 2017; Riising et al., 2014; Shan et al., 2017). In contrast, primed PSCs are sensitive to the removal of Polycomb proteins (Collinson et al., 2016; Moody et al., 2017; Shan et al., 2017; Wang et al., 2018). These observations collectively imply that the stable transition from a naïve to a primed state of pluripotency requires the reconfiguration of DNA interactions to provide a coordinated set of ‘poised’ regulatory signals to control promoter priming.

A clear difference between human pluripotent states that we observed was in enhancer interactivity and activity state, and these differences were associated with the widespread reorganisation of transcription factor binding. The shared factors OCT4, SOX2 and NANOG bound predominantly to active enhancers in both pluripotent states. There was, however, a remarkable lack of overlap in OSN-occupied enhancers between naïve and primed PSCs. This finding suggests that there is a substantial reorganisation of gene regulatory elements between human pluripotent states. In mouse PSCs, enhancer activation and OSN occupancy are also dynamic between pluripotent states (Buecker et al., 2014; Factor et al., 2014; Galonska et al., 2015) and these processes are modulated by the presence of other state-specific transcription factors such as ESRRB in naïve cells and OTX2 and GRHL2 in primed cells (Atlasi et al., 2019; Buecker et al., 2014; Chen et al., 2018; Yang et al., 2014). Interestingly, *ESRRB* is not expressed in naïve human PSCs nor in human epiblast cells in vivo (Blakeley et al., 2015; Petropoulos et al., 2016; Stirparo et al., 2018), suggesting that in some instances the control of enhancer activity differs between mouse and human pluripotent states and that alternative factors are involved in human cells. One such factor is TFAP2C, which facilitates the opening of a subset of naïve-specific enhancers in human PSCs but only has a modest role in mouse PSCs (Pastor et al., 2018). The emerging picture is that substantial rewiring of enhancer activity and interactions occurs during pluripotent state transitions and this is driven by a combination of common and species-specific transcription factors. An added complexity is that the cooperative behaviour of multiple transcription factors seems to be context-dependent in terms of which enhancers are targeted, and this complexity will need to be unravelled through precise functional analyses. Our results will transform the interpretation of these experiments by comprehensively identifying the target gene promoters for enhancers and their dynamic alteration in human pluripotent states.

The *Canvas* network approach that we developed here represents patterns of interactions (edges) between genomic regions (nodes), with added edge weights and node values that can convey user-defined information such as cell-type specificity, the chromatin or transcriptional state of each region, or the interaction strength. We chose to visualise the combined network using a force-directed layout (Jacomy et al., 2014). Other visualisation algorithms that we tested were less suitable because they generated network graphs that lacked informative structure or were incompatible with large-scale data sets. The final, constructed networks revealed prominent features of genome organisation at multiple scales such as compartments, TADs and chromatin loops. *Canvas*, therefore, provides a powerful and intuitive data exploration method to understand the relationships behind the emergent connectivity patterns. Our network-scale approach could be applied to other interaction datasets and we provide our scripts and documentation to facilitate this. Future iterations of the network graph could take into account other features of spatial organisation such as chromosome territories and positioning, and lamin-associated domains, to instruct the layout of nodes and edges within the network. Another area of future development could be to adapt the existing network to make predictions about how the set of interactions formed or how the interactions might change upon perturbation or cell state changes. Such theory-based models typically require a higher level of abstraction away from the experimental findings, but are able to make network-dependent predictions about the mechanisms and dynamics of the interaction model.

Our study provides an invaluable resource to study the complex interplay between transcription factors, chromatin state and 3D genome organisation in controlling cell state. By visualising chromatin interactions as a network, we uncover the substantial and multilayered differences between two human pluripotent states. This approach could be applied to other high-resolution interaction datasets to uncover topological changes that underpin cell state transitions during development and disease.

## ACKNOWLEDGMENTS

We thank Steven Wingett, Felix Krueger and Simon Andrews from Babraham Bioinformatics for sequencing QC and mapping, and assistance with bioinformatic analysis; and Kristina Tabbada and Clare Murnane of the Babraham Institute Next Generation Sequencing Facility. We are also grateful to Mikhail Spivakov and Jonathan Cairns for advice about CHiCAGO and Peter Fraser for support and helpful discussions. Work in our laboratories is supported by grants from the BBSRC (BBS/E/B/000C0421, BBS/E/B/000C0422, BB/J004480/1, Core Capability Grant); A.J.C. was supported by an MRC DTG Studentship (MR/J003808/1). S.S. was supported by a UKRI MRC Rutherford Fund Fellowship (MR/T016787/1). C.V. was supported by an ERC AdG to Peter Fraser (DEVOCHROMO).

## AUTHOR CONTRIBUTIONS

Conceptualisation, P.C., A.J.C., S.S., A.C. and P.J.R.-G; Methodology, P.C. and A.J.C; Software, P.C; Validation, P.C., A.J.C., C.K., C.V., S.S., A.C. and P.J.R.-G; Formal analysis, P.C., A.J.C., C.K. and C.V; Investigation, P.C., A.J.C. and S.S; Resources, P.C., A.J.C., S.S., A.C. and P.J.R.-G; Data Curation, P.C., A.J.C., C.K. and C.V; Writing - Original Draft, P.C., A.J.C. C.K., C.V. and P.J.R.-G; Writing - Review and Editing, P.C., A.J.C. C.K., C.V. S.S., A.C. and P.J.R.-G; Visualization, P.C., A.J.C. C.K. and C.V; Supervision, S.S., A.C. and P.J.R.-G; Project Administration, P.C., A.J.C., S.S., A.C. and P.J.R.-G; Funding Acquisition, A.C. and P.J.R.-G.

## METHODS

### Cell lines

WA09/H9 NK2 naïve and primed PSCs were kindly provided by Dr. Austin Smith (Takashima et al., 2014) with permission from WiCell. All PSCs were cultured in 5% O_2_, 5% CO_2_ at 37°C.

### Cell culture

Naïve PSCs were maintained as previously described (Takashima et al., 2014) in a 1:1 mixture of DMEM/F12 and Neurobasal, 0.5x N2 supplement, 0.5x B27 supplement, 1x nonessential amino acids, 2mM L-Glutamine, 1x Penicillin/Streptomycin (all from ThermoFisher Scientific), 0.1mM β-mercaptoethanol (Sigma-Aldrich), 1 μM PD0325901, 1 μM CHIR99021, 20ng/ml human LIF (all from WT-MRC Cambridge Stem Cell Institute) and 2 μM Gö6983 (PKCi; Tocris) on Matrigel–coated plates (Corning). Primed PSCs were maintained on Vitronectin-coated plates (0.5 μg/cm^2^; ThermoFisher Scientific) in TeSR-E8 medium (StemCell Technologies).

### Hi-C

Hi-C libraries were generated essentially as described (Schoenfelder et al., 2015b) with modifications detailed below. Approximately 35 million naïve or primed PSCs were fixed in 2% formaldehyde (Agar Scientific) for 10 min, after which the reaction was quenched with ice-cold glycine (Sigma; 0.125M final concentration). Cells were collected by centrifugation (400 x g for 10 min at 4°C), and washed once with 50 mL PBS pH 7.4 (Gibco). After another centrifugation step (400 x g for 10 min at 4°C), the supernatant was completely removed and the cell pellets were immediately frozen in liquid nitrogen and stored at −80°C. After thawing, the cell pellets were incubated in 50 mL ice-cold lysis buffer (10 mM Tris-HCl pH 8, 10 mM NaCl, 0.2% Igepal CA-630, cOmplete EDTA-free protease inhibitor cocktail (Roche)) for 30 min on ice. After centrifugation to pellet the cell nuclei (650 x g for 5 min at 4°C), nuclei were washed once with 1.25 x NEBuffer 2 (NEB). The nuclei were then resuspended in 1.25 x NEBuffer 2, SDS (10% stock; Promega) was added (0.3% final concentration) and the nuclei were incubated at 37°C for one hour with agitation (950 rpm). Triton X-100 (Sigma) was added to a final concentration of 1.7% and the nuclei were incubated at 37°C for one hour with agitation (950 rpm). Restriction digest was performed overnight at 37°C with agitation (950 rpm) with *HindIII* (NEB; 1500 units per 7 million cells). Using biotin-14-dATP (Life Technologies), dCTP, dGTP and dTTP (Life Technologies; all at a final concentration of 30 μM), the *HindIII* restriction sites were then filled in with Klenow (NEB) for 75 minutes at 37°C, followed by ligation for 4 hours at 16°C (50 units T4 DNA ligase (Life Technologies) per 7 million cells starting material) in a total volume of 5.5 mL ligation buffer (50 mM Tris-HCl, 10 mM MgCl2, 1 mM ATP, 10 mM DTT, 100 μg/mL BSA) per 7 million cells starting material. After ligation, crosslinking was reversed by incubation with Proteinase K (Roche; 65 μl of 10mg/mL per 7 million cells starting material) at 65°C overnight. An additional Proteinase K incubation (65 μl of 10mg/mL per 7 million cells starting material) at 65°C for two hours was followed by RNase A (Roche; 15 μl of 10mg/mL per 7 million cells starting material) treatment and two sequential phenol/chloroform (Sigma) extractions. After DNA precipitation (sodium acetate 3M pH 5.2 (1/10 volume) and ethanol (2.5 x volumes)) overnight at −20°C, the DNA was spun down (centrifugation 3200 x g for 30 min at 4°C). The pellets were resuspended in 400 μl TLE (10 mM Tris-HCl pH 8.0; 0.1 mM EDTA), and transferred to 1.5 mL eppendorf tubes. After another phenol/chloroform (Sigma) extraction and DNA precipitation overnight at −20°C, the pellets were washed three times with 70% ethanol, and the DNA concentration was determined using Quant-iT Pico Green (Life Technologies). For quality control, candidate 3C interactions were assayed by PCR, and the efficiency of biotin incorporation was assayed by amplifying a 3C ligation product, followed by digest with *HindIII* or *NheI*.

To remove biotin from non-ligated fragment ends, 40 μg of Hi-C library DNA were incubated with T4 DNA polymerase (NEB) for 4 hours at 20°C, followed by phenol/chloroform purification and DNA precipitation overnight at −20°C. After one wash with 70% ethanol, sonication was carried out to generate DNA fragments with a size peak around 400 bp (Covaris E220 settings: duty factor: 10%; peak incident power: 140W; cycles per burst: 200; time: 55 seconds). After end repair (T4 DNA polymerase, T4 DNA polynucleotide kinase, Klenow (all NEB) in the presence of dNTPs in ligation buffer (NEB)) for 30 min at room temperature, the DNA was purified (Qiagen PCR purification kit). dATP was added with Klenow exo- (NEB) for 30 min at 37°C, after which the enzyme was heat-inactivated (20 min at 65°C). A double size selection using AMPure XP beads (Beckman Coulter) was performed: first, the ratio of AMPure XP beads solution volume to DNA sample volume was adjusted to 0.6:1. After incubation for 15 min at room temperature, the sample was transferred to a magnetic separator (DynaMag-2 magnet; Life Technologies), and the supernatant was transferred to a new eppendorf tube, while the beads were discarded. The ratio of AMPure XP beads solution volume to DNA sample volume was then adjusted to 0.9:1 final. After incubation for 15 min at room temperature, the sample was transferred to a magnet (DynaMag-2 magnet; Life Technologies). Following two washes with 70% ethanol, the DNA was eluted in 100 μl of TLE (10 mM Tris-HCl pH 8.0; 0.1 mM EDTA). Biotinylated ligation products were isolated using MyOne Streptavidin C1 Dynabeads (Life Technologies) on a DynaMag-2 magnet (Life Technologies) in binding buffer (5 mM Tris pH8, 0.5 mM EDTA, 1 M NaCl) for 30 min at room temperature. After two washes in binding buffer and one wash in ligation buffer (NEB), PE adapters (Illumina) were ligated onto Hi-C ligation products bound to streptavidin beads for 2 hours at room temperature (T4 DNA ligase NEB, in ligation buffer, slowly rotating). After washing twice with wash buffer (5 mM Tris, 0.5 mM EDTA, 1 M NaCl, 0.05% Tween-20) and then once with binding buffer, the DNA-bound beads were resuspended in a final volume of 90 μl NEBuffer 2. Bead-bound Hi-C DNA was amplified with 7 PCR amplification cycles using PE PCR 1.0 and PE PCR 2.0 primers (Illumina). After PCR amplification, the Hi-C libraries were purified with AMPure XP beads (Beckman Coulter). The concentration of the Hi-C libraries was determined by Bioanalyzer profiles (Agilent Technologies), and the Hi-C libraries were paired-end sequenced (HiSeq 2500, Illumina).

### Promoter Capture Hi-C

Promoter Capture Hi-C libraries were generated essentially as described (Schoenfelder et al., 2015b) with modifications detailed below. 500 ng of Hi-C library DNA was resuspended in 3.6 μl water, and hybridization blockers (Agilent Technologies; hybridization blockers 1 and 2, and custom hybridization blocker) were added to the Hi-C DNA. Hybridization buffers and the custom-made RNA capture bait system (Agilent Technologies; designed as previously described (Mifsud et al., 2015): 37,608 individual biotinylated RNAs targeting the ends of 22,076 promoter-containing human *HindIII* restriction fragments) were prepared according to the manufacturer’s instructions (SureSelect Target Enrichment, Agilent Technologies). The Hi-C library DNA was denatured for 5 min at 95°C, and then incubated with hybridization buffer and the RNA capture bait system at 65°C for 24 hours (all incubation steps in a MJ Research PTC-200 PCR machine). After hybridization, 60 μl of MyOne Streptavidin T1 Dynabeads (Life Technologies) were washed three times with 200 μl binding buffer (SureSelect Target Enrichment, Agilent Technologies), before incubation with the Hi-C DNA/RNA capture bait mixture with 200 μl binding buffer for 30 min at room temperature, slowly rotating. Hi-C DNA bound to capture RNA was isolated using a DynaMag-2 magnet (Life Technologies). Washes (15 min in 500 μl wash buffer I at room temperature, followed by three 10 min incubations in 500 μl wash buffer II at 65°C) were performed according to the SureSelect Target enrichment protocol (Agilent Technologies). After the final wash, the beads were resuspended in 300 μl NEBuffer 2, isolated on a DynaMag-2 magnet, and then resuspended in a final volume of 30 μl NEBuffer 2. After a post-capture PCR (four amplification cycles using Illumina PE PCR 1.0 and PE PCR 2.0 primers; 13 to 15 individual PCR reactions), the Promoter CHi-C libraries were purified with AMPure XP beads (Beckman Coulter). The concentration of the Promoter CHi-C libraries was determined by Bioanalyzer profiles (Agilent Technologies), and the Promoter CHi-C libraries were paired-end sequenced (HiSeq 2500, Illumina).

### Hi-C analysis

HiCUP (Wingett et al., 2015) was used to map and filter di-tags to human genome build GRCh38. The aligned Hi-C data were normalized using HOMER v4.7 (Heinz et al., 2010) and Juicer tools v1.8.9 (Durand et al., 2016). Using binned Hi-C data, we computed the coverage- and distance-related background in the Hi-C data employing matrix balancing algorithms at 25 kb and 250 kb (iterative correction by HOMER) and a 5 kb to 2.5 Mb (Knight–Ruiz balancing by Juicer tools) range of resolutions. We compared global organisation by plotting the log10 frequency of cis-chromosomal contacts in the raw data at various genomic distances on the log10 scale.

The spatial compartment signal was computed as the first principle component of the distance- and coverage-corrected interaction profile correlation matrix at 250 kb resolution (Lieberman-Aiden et al., 2009), with positive values aligned with H3K4me3 ChIP-seq data. Using ‘A’ and ‘B’ compartment bins on the autosomal chromosomes defined by the positive and negative compartment signal values, we computed a compartmentalisation score as the log2 enrichment of contacts between the A and B compartments using the trans- or the long-cis- (>10Mb) chromosomal contacts. To compute contact enrichment matrices, we defined 50 equally sized compartment groups of increasing A compartment association using their compartment signal values in the corresponding dataset (with group 1 being the most ‘B’ compartment-like and group 50 being the most ‘A’ compartment-like) and computed pairwise log2 contact enrichments between these 50 groups. Expected contact frequencies were computed using +10 Mb shifted compartment signal. For compartment switching between naïve and primed PSCs we only considered compartments with a score >5 and <-5.

TADs were identified based on directionality indices of Hi-C interactions (Dixon et al., 2012), using HOMER with minDelta=2 and other parameters kept at their default values. This resulted in 3,124 TADs for naïve PSCs and 2,917 TADs for primed PSCs.

Hi-C peaks in Figure 4 were identified using HiCCUPS v1.8.8 (Rao et al., 2014) with the following parameters: --ignore_sparsity -k KR -f 0.1 -r 250000 -d 750000 -i 8 -p 4. The peaks identified on chromosome 5 in primed PSCs were used for both naïve and primed aggregate peak analysis with the following parameters: -r 250000 -c chr5 -n 30 -w 10 using Juicer tools v1.8.9 (Durand et al., 2016).

### Promoter Capture Hi-C analysis

PCHi-C data were mapped and filtered using HiCUP (Wingett et al., 2015) with the GCRh38 human genome build. CHiCAGO (Cairns et al., 2016) was used to define significant promoter interactions at the level of individual *HindIII* fragments. Two biological replicates for each cell type were normalised and combined as part of the CHiCAGO pipeline. CHiCAGO interaction scores correspond to –log-transformed, weighted p-values for each fragment read pair. A CHiCAGO interaction score of 5 or above was considered significant based on previous empirical observations (Cairns et al., 2016; Freire-Pritchett et al., 2017). The network graph was constructed using promoter interactions with scores of 5 or above. For Figures 6 and 7, to avoid removing interactions that were close to the applied threshold, interactions with scores between 3 and 5 were included in the analysis if they scored between 5 and 7 in the other cell type.

### Network construction and analysis

PCHi-C interactions with CHiCAGO scores of 5 or more were used to generate the network graph using custom scripts in R and Python. Promoter regions were defined as 1 kb upstream of the TSS based on the Ensembl v85 gene model annotation and assigned to *HindIII* fragments. Interaction networks were constructed using the igraph v1.2.1 R package (Csardi and Nepusz, 2006) where each *HindIII* fragment represents a node and each significant interaction represents an edge. The edges of both naïve and primed PSCs were combined into a single network with a common layout. The shared anchor points allow examination of interactivity of shared genomic regions. For Figure 2, community detection was performed using the multi-level optimisation algorithm (Blondel et al., 2008) implemented within igraph. Subnetworks with a modularity score of 0.7 or above were split into individual communities. The coordinates of *HindIII* fragments with the assigned communities were compared to TAD coordinates obtained from Hi-C analysis using the GenomicRanges R package v1.30.3 (Lawrence et al., 2013). Network visualisation was performed within Gephi v0.9.2 (Bastian M., Heymann S., Jacomy M., 2009) using the ForceAtlas2 layout algorithm (Jacomy et al., 2014). For Figure 4B, the multi-dimensional scaling (MDS) layout within Gephi, developed by Wouter Spekkink (http://www.wouterspekkink.org), was used to obtain linear genomic distance representative layouts of individual subnetworks.

For Figure S5D, interaction distances >1.5 times the interquartile range were classified as outliers and not plotted.

### Chromatin immunoprecipitation

All buffers were pre-chilled to 4°C with cOmplete EDTA-free protease inhibitor (Roche) freshly added. 15 million cells per ChIP were treated with Accutase (ThermoFisher Scientific) and collected in a 50ml conical tube, followed by a 300 x g 5min spin at 4°C. The pellets were resuspended in PBS. Cells were cross-linked with 2µM DSG (Sigma) for 45min at 22°C and then with 1% methanol-free formaldehyde (Agar Scientific) at a cell density of 10^8^ cells in 45ml media for 12.5min at 22°C. Fixation was stopped with the addition of glycine at a final concentration of 125mM and incubation for 5min at 22°C. After two PBS washes, cells were resuspended in Wash buffer 1 (10mM Hepes pH 7.5; 10mM EDTA; 0.5mM EGTA; 0.75% Triton X-100) and incubated for 10min at 4°C. After spinning at 3200 x g for 5min at 4°C, nuclei were resuspended in 10ml Wash buffer 2 (10mM Hepes pH 7.5; 200mM NaCl; 1mM EDTA; 0.5mM EGTA) and incubated for 10min at 4°C. Another 3200 x g 5min spin at 4°C was performed followed by resuspension in 1ml of freshly made Lysis/sonication buffer (150mM NaCl; 25mM Tris pH 7.5; 5mM EDTA; 0.1% Triton X-100; 1% SDS; freshly dissolved 0.5% Sodium deoxycholate) per 12 million cells. Lysis was performed on ice for 30min, followed by sonication for 15 seconds on, 30 seconds off (Microson ultrasonic cell disruptor XL Misonix; output setting 4; 10-11 W) for 20 cycles to obtain fragments with a size of 200-500bp. Fragmented chromatin was spun down at 10,000 x g for 15min at 4°C, and the supernatant was transferred to a new tube and diluted 1:10 with ChIP dilution buffer (150 mM NaCl; 25mM Tris pH 7.5; 5mM EDTA; 1% Triton X-100; 0.1% SDS; 0.5% Sodium deoxycholate). 500µl was taken for the input and the remaining diluted supernatant was incubated with 5µg of antibody overnight at 4°C. Magnetic protein A (120µl per IP) or protein G (180µl per IP) Dynabeads (both from Invitrogen) were washed with Wash buffer A (50mM Tris pH 8; 150mM NaCl; 0.1% SDS; 0.5% Sodium deoxycholate; 1% NP40; 1mM EDTA) and blocked for 1 hour at 4°C with yeast tRNA (Invitrogen) and BSA (NEB). The pre-blocked beads were added to the antibody-bound chromatin and incubated for 7-8 hours at 4°C. Subsequently, the magnetic beads with the bound antibody-chromatin-complex were rinsed once with Wash buffer A, washed twice with Wash buffer A, washed once with Wash buffer B (50mM Tris pH 8.0; 500mM NaCl; 0.1% SDS; 0.5% Sodium deoxycholate; 1% NP40; 1mM EDTA), washed once with Wash buffer C (50mM Tris pH 8; 250mM LiCl; 0.5% Sodium deoxycholate; 1% Igepal CA-630; 1mM EDTA) and rinsed with 1x TE buffer (10mM Tris pH 8; 1mM EDTA). Chromatin was eluted off the beads with 450µl of Elution buffer (1% SDS; 0.1M NaHCO_3_). Additionally, 11µl Proteinase K (20mg/ml) and 5µl RNase A (10mg/ml) were added (including to the input) and incubated at 37°C for 2 hours, followed by an overnight incubation at 65°C to reverse the crosslink. DNA was purified using AMPure XP beads (Beckman Coulter) and eluted in 40µl water. DNA was quantified using the Qubit fluorometer dsDNA HS assay kit (ThermoFisher Scientific). Libraries were prepared using the NEBNext Ultra II DNA library prep kit for Illumina (NEB) using the manufacturer’s protocol.

### Genome browser tracks

Normalised bigwig files for genome browser visualisation were produced using Deeptools (Ramírez et al., 2016). For ChIP-seq samples, the BAM files were normalised with the reads per genomic content (RPGC) method, ignoring chrY and chrMT. A 10 bp bin size with a 200 bp read extension was chosen with a bigwig file output. The experimental inputs were subtracted from each sample. For RNA-seq tracks, the BAM files were normalised using the DESeq2 scaling factor (Naïve - 0.34576529; Primed - 1.873937595), with a default bin size of 50 bp. Genome browser tracks were visualised using the WashU Epigenome Browser v48.2.0+ (Zhou et al., 2011, 2013, 2015).

### Quantification and statistical analysis

#### ChIP-sequencing analysis

Reads were trimmed using Trim Galore and mapped to human genome GRCh38 using Bowtie2 (Langmead and Salzberg, 2012). All analyses were performed using SeqMonk and R. ChIP-seq peaks were called using MACS2 (Zhang et al., 2008) with parameters q<10^-9^ for all histone modification samples except for H3K4me1 for which the cutoff used was q<10^-7^. For quantitation, read lengths were extended to 300 bp and regions of coverage outliers were excluded. Quantitations (log2 RPM) for all analysed regions and histone marks are provided in the OSF (ChIP_Quantiation). OCT4, NANOG and SOX2 peaks were called using a SeqMonk implementation of MACS (Zhang et al., 2008) with parameters p<10^-5^, sonicated fragment size = 300. Peaks were filtered by signal intensity, retaining only peaks that overlap with at least one 500 bp window in which log2 RPM > 0. Regions of OCT4, NANOG and SOX2 peaks were combined and merged if closer than 100 bp. The resulting list of regions was filtered for those that overlap with MACS peaks for all three factors and called OSN peaks. Control regions are 10000 randomly selected 1.2 kb windows (approximate average peak size). To assign one ChromHMM state per peak, the location of the peak centre was used.

In Figures 6A to 6H, enhancers and super-enhancers were annotated using ROSE (Lovén et al., 2013; Whyte et al., 2013) with H3K27ac peaks called using MACS2 with parameters q<10^-9^. A stitching distance of 1.5 kb was chosen based on the bimodal distribution of the distance to the nearest peak.

#### Chromatin state annotation - ChromHMM

Chromatin state analysis was performed using ChromHMM (Ernst and Kellis, 2012). Trim Galore quality trimmed and Bowtie2 aligned (GRCh38) BAM files were binarized using the BinarizeBam command with default 200 bp bin settings. Naïve and primed PSCs were stacked to provide a single-genome annotation with the inclusion of ChIP-seq input samples as an additional feature. Model learning was performed on a range of states with 16 being selected as the final number. These categories were reduced to seven states that were more biologically relevant (Active – H3K27ac, H3K4me3, H3K4me1; Polycomb-repressed – H3K27me3; Bivalent – H3K27me3, H3K4me3, H3K4me1; Heterochromatin repressed – H3K9me3; Unclassified - H3K27me3, H3K4me3, H3K4me1, H3K9me3; Background – low emission probability levels in all samples).

To assign a state to each *HindIII* fragment, the overlap of the seven states with each *HindIII* restriction fragment was determined and reduced to a single final chromatin state based on the following rules: any single state superseded the background state; the bivalent state superseded the Polycomb-repressed state, and a mixture of multiple states was labelled as mixed. Genomic features relating to ChromHMM states were used in Figures 6I, 6J, 7, S6E and S6F and their definitions are summarised in Table S3.

#### RNA-Seq analysis

RNA-sequencing reads from (Takashima et al., 2014) were trimmed using Trim Galore 0.3.8 using default parameters to remove the standard Illumina adaptor sequence. Reads were mapped to the human GRCh38 genome assembly using HISAT 2.0.5 (Kim et al., 2015) guided by the gene models from the Ensembl v85 release. Samtools (Li et al., 2009) was used to convert to BAM files that were imported to Seqmonk. Raw read counts per transcript were calculated using the RNA-sequencing quantitation pipeline on the Ensembl v85 gene set using directional counts. Differentially expressed genes were determined using DESeq2 (Love et al., 2014). Log2(FPKM) normalised values were generated with Seqmonk.

#### DNA methylation analysis

Whole-genome bisulfite sequencing data from (Takashima et al., 2014) was trimmed using Trim Galore 0.3.8, aligned with Bismark v0.18.2 (Krueger and Andrews, 2011) and analysed in SeqMonk. Methylation is given as percentage of methylated cytosine calls over all observations for each enhancer or background region.

#### Gene ontology analysis

Gene ontology analysis of protein-coding genes within the H3K27me3-associated interaction network in primed PSCs was performed using Enrichr (Chen et al., 2013; Kuleshov et al., 2016).

#### Motif enrichment

To find differentially enriched motifs between naïve and primed specific OSN binding sites, sequences 250 bp up- and downstream of the peak centre were repeat masked (RepeatMasker, A.F.A. Smit, R.Hubley & P.Green, version open-4.0.9, default parameters). 500 OSN peaks were randomly selected per category and analysed using AME (version 5.0.5, (McLeay and Bailey, 2010)) with the following parameters (‘naïve over primed’): ame --verbose 1 --oc. --scoring avg --method fisher --hit-lo-fraction 0.25 --evalue-report-threshold 10.0 --control 500_random_primed_specific_OSN.txt.fa 500_random_naive_specific_OSN.txt.fa db/JASPAR/JASPAR2018_CORE_vertebrates_non-redundant.meme. For enrichment ‘primed over naïve’ the two input files were swapped. Resulting lists of enriched motifs were filtered for expression of the respective binding factor in at least one of the two PSC states (log2 RPKM > 0). A FIMO search (Grant et al., 2011) using the sequences of all selected OSN regions as a background model was then performed to determine whether an individual OSN peak contained the motif or not.

## Data and software availability

All raw and processed data and custom scripts will be made available upon publication.

## SUPPLEMENTAL FIGURES

**Figure S1.**
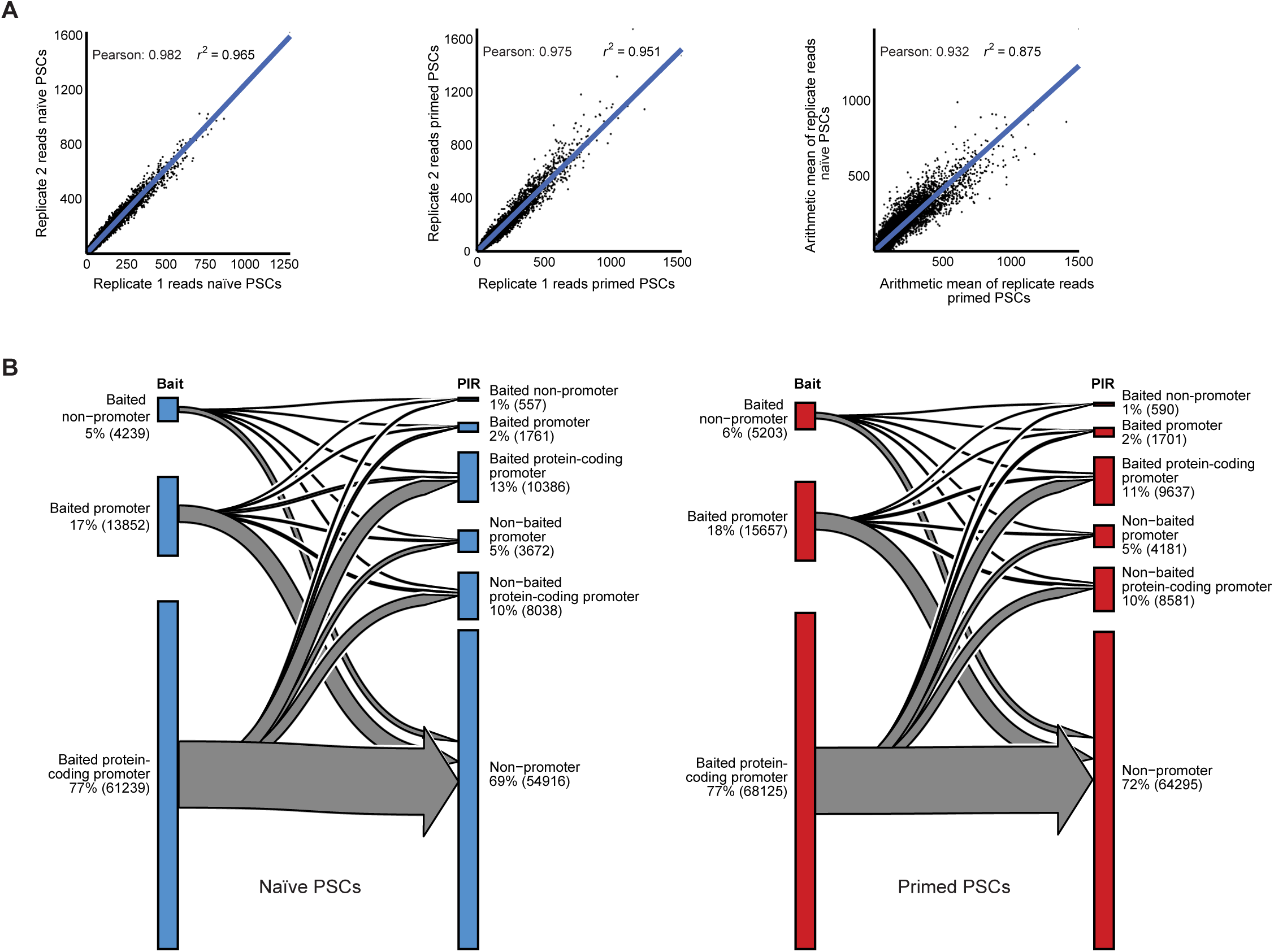
Characterisation of PCHi-C replicates and interacting regions. (A) Scatter plots show pairwise comparisons between individual PCHi-C replicates (naïve, left; primed, centre) and between the mean of replicate reads in naïve and primed PSCs (right). Correlation values are shown. (B) Sankey plots describe the genomic features of interacting regions in naïve PSCs (left) and primed PSCs (right) that were detected by PCHi-C. Listed within the ‘bait’ category are the number and percentage of protein-coding promoters, non-protein-coding promoters and non-promoter regions. The ‘PIR’ category describes the promoter-interacting regions. As expected, the most frequent type of interaction in both cell types was between baited protein-coding promoters and non-promoter regions.

**Figure S2.**
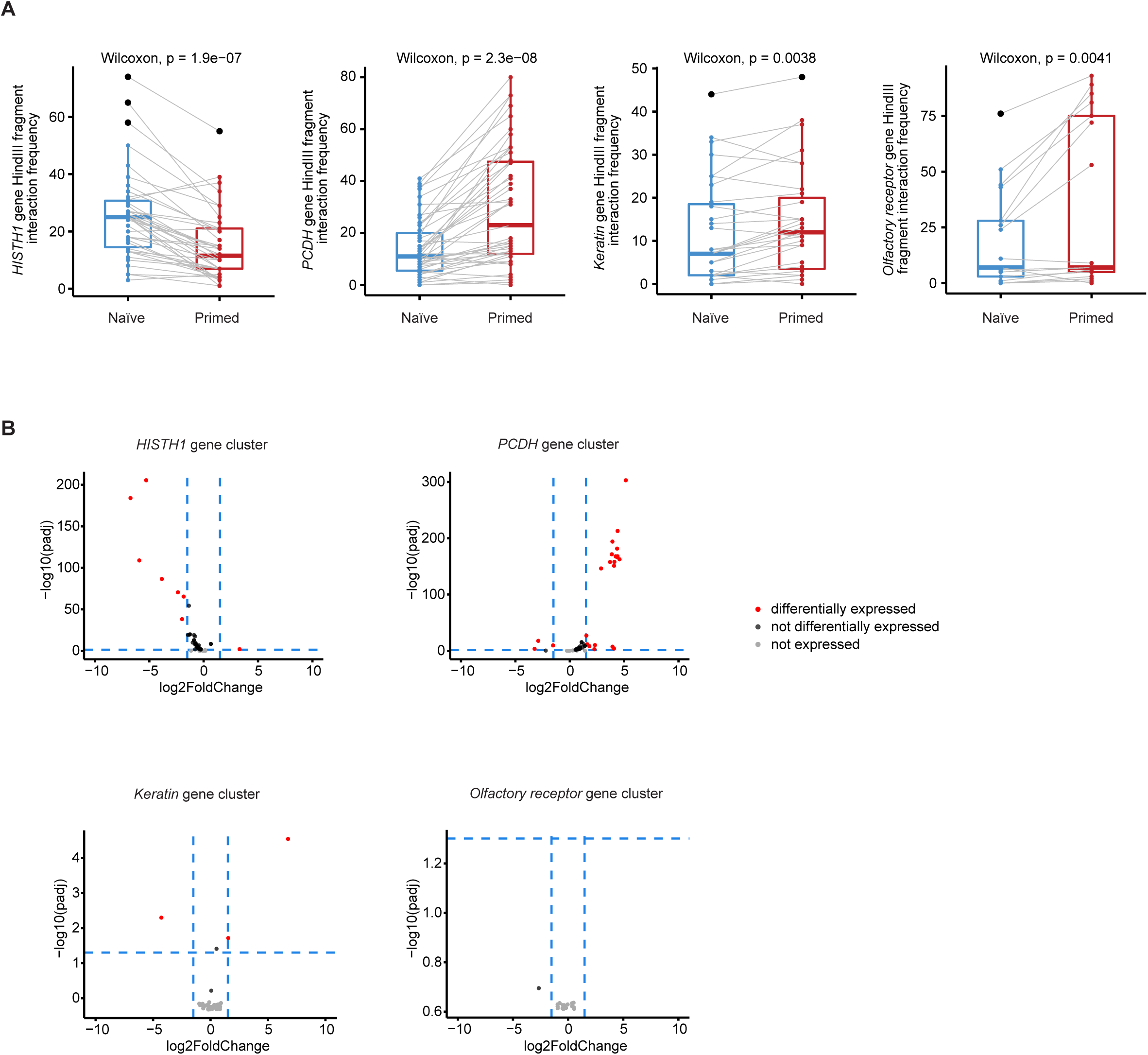
Changes in promoter-interaction frequency and transcriptional levels in four gene clusters. (A) Box plots show the number of significant interactions at *HindIII* fragments located within the *HISTH1*, *PCDH*, *KRT* and Olfactory gene clusters. (B) Volcano plots show the transcriptional changes between naïve and primed PSCs for genes within the *HISTH1*, *PCDH*, *KRT* and Olfactory gene clusters. Each dot represents a different gene. Genes coloured in red are differentially expressed between naïve and primed PSCs (log10 fold change >1.5 or > −1.5 and with an adjusted p-value < 0.05). Other categories shown include genes that are not differentially expressed (log10 fold change <1.5 < −1.5 and/or with an adjusted p-value > 0.05) or genes that are not expressed (0 read counts in RNA-seq data).

**Figure S3.**
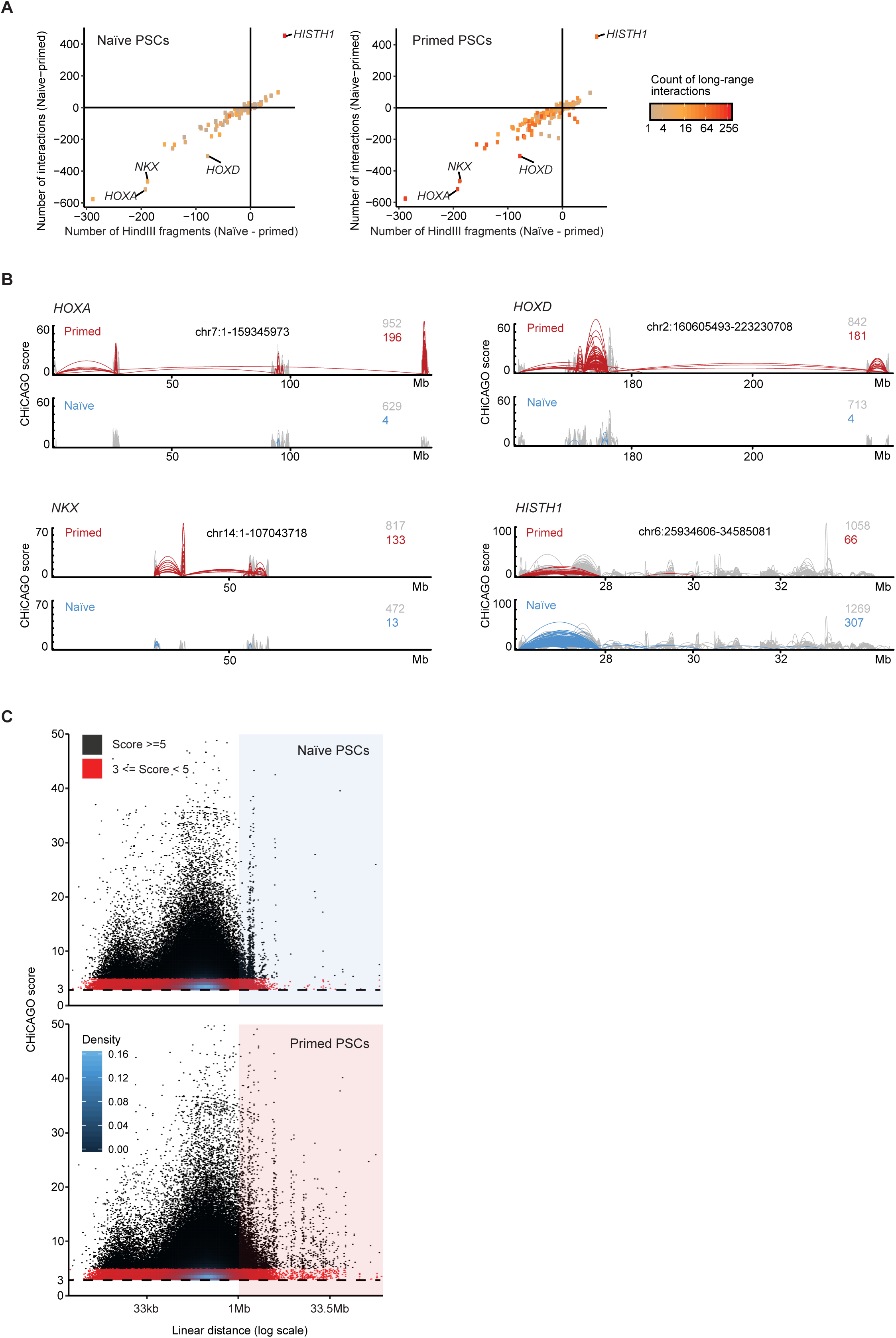
Long-range promoter interactions create large sub-networks in primed PSCs. (A) Scatter plots show the number of interactions (edges) and the number of interacting *HindIII* fragments (nodes) for each sub-network in naïve and primed PSCs. The lower-left quadrant contains larger sub-networks in primed PSCs, and the upper-right quadrant contains larger sub-networks in naïve PSCs. The *HOXA*, *HOXD*, *NKX* and *HISTH1* sub-networks are highlighted. Sub-networks are coloured according to their number of long-range promoter interactions. Note the increased number of long-range promoter interactions within most sub-networks in primed (right) compared to naïve (left) PSCs. (B) Genome browser tracks show the PCHi-C interactions and CHiCAGO scores in naïve and primed PSCs for the *HOXA*, *HOXD*, *NKX* and *HISTH1* sub-networks. (C) Dot plots show that the high number of long-range promoter interactions in primed PSCs is independent of the applied CHiCAGO threshold. Each dot represents a PCHi-C interaction, positioned according to the linear genomic distance of the interaction (x-axis) and the assigned CHiCAGO score (y-axis). Black dots show the interactions obtained when applying a CHiCAGO score of >5 (the threshold used for constructing the network graph) and red dots show the interactions when using a relaxed CHiCAGO score of between 3 and 5. Primed PSCs have more long-range promoter interactions (shaded area; defined as >1Mb) compared to naïve PSCs when either CHiCAGO threshold score is applied.

**Figure S4.**
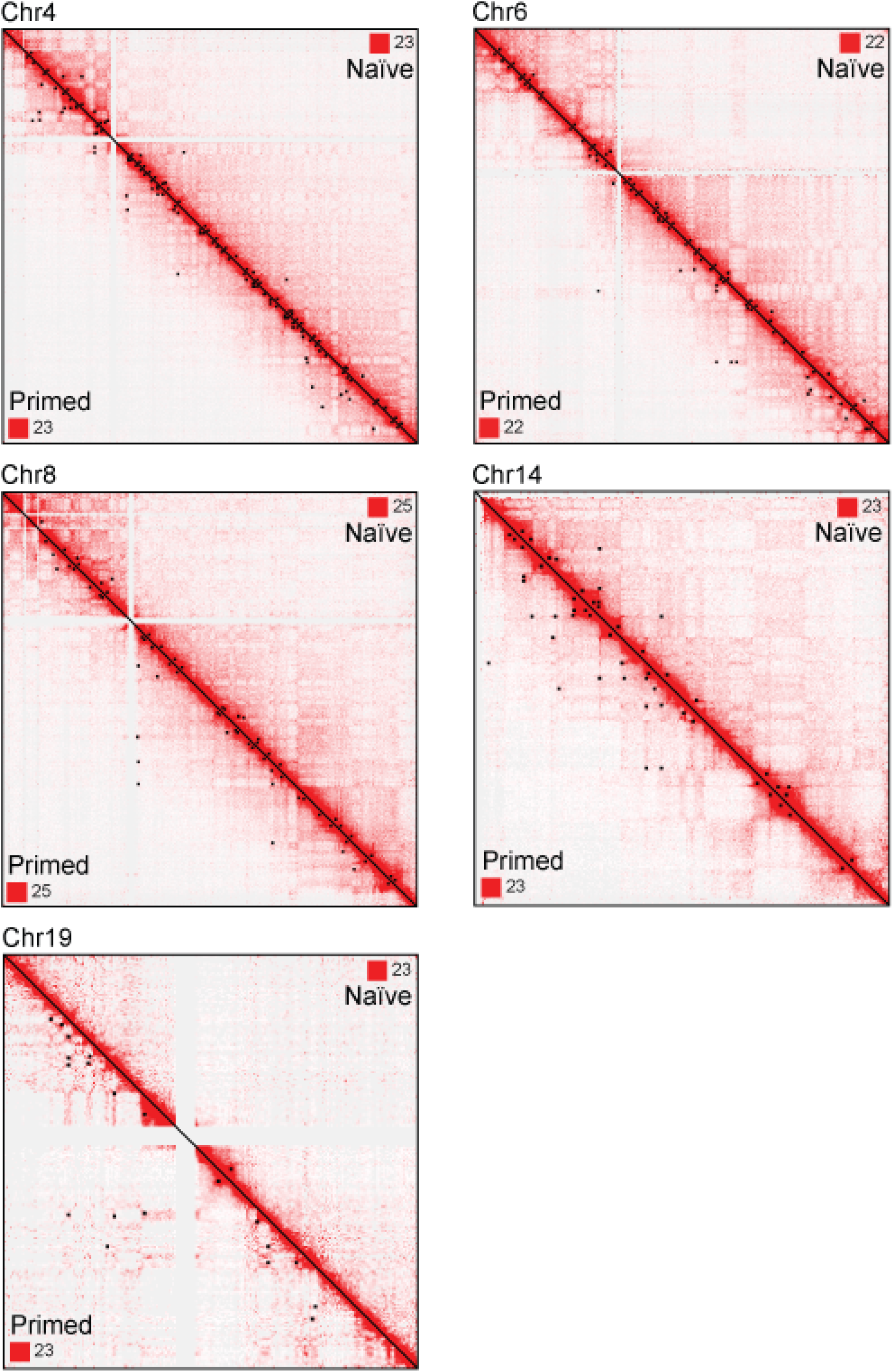
Additional examples show the higher number of long-range chromatin interactions in primed compared to naïve PSCs. Five examples from different chromosomes of Hi-C interaction matrices at a resolution of 250 kb with Knight-Ruiz (KR) normalisation. In each contact matrix, naïve PSCs are shown in the upper right and primed PSCs are shown in the lower left. Areas of contact enrichment were defined separately for naïve and primed PSCs using HiCCUPS analysis of Hi-C data and each cell type-specific set of chromatin interactions are highlighted as a black square on their respective heatmaps. The numbers in each corner indicate the maximum intensity values for the matrix. Chovanec Figure S4

**Figure S5.**
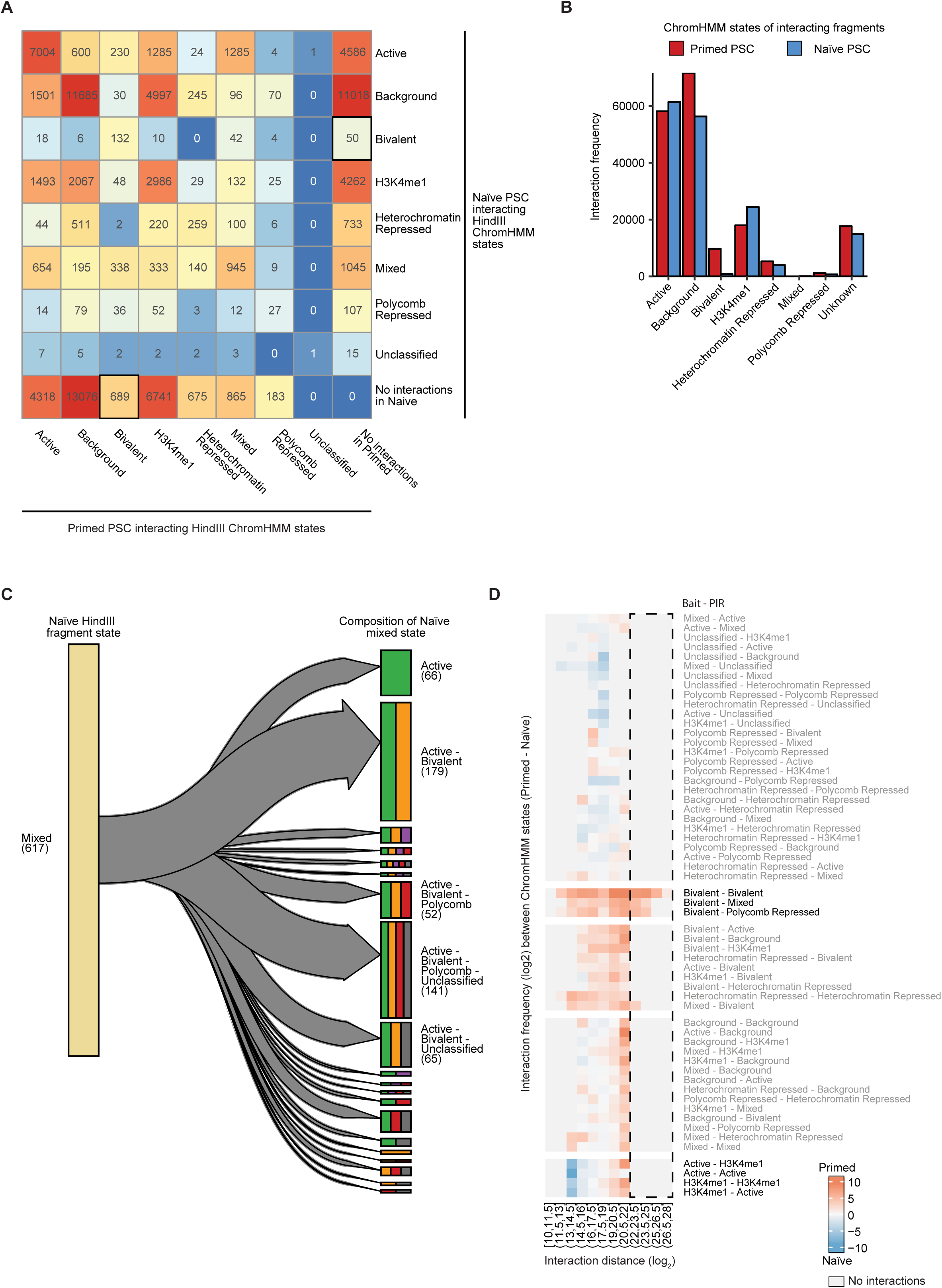
Assigning ChromHMM states to chromatin interacting regions. (A) Heatmap shows the number of interacting *HindIII* fragments that were assigned to each of the ChromHMM-defined chromatin states in both cell types. Note the higher number of bivalently-marked (H3K27me3 and H3K4me1/3) interacting regions in primed compared to naïve PSCs. (B) Chart shows the total number of interacting *HindIII* fragments for each of the ChromHMM states. (C) Sankey plot reveals the chromatin state composition for the 617 *HindIII* interacting regions in naïve PSCs that were defined by ChromHMM as being in a ‘mixed’ chromatin state. The left column shows the 617 mixed chromatin state regions and the right column shows the breakdown of individual chromatin states for each of the regions, represented by the different colour bars. Approximately half of the mixed state *HindIII* fragments contain signatures of active (H3K4me3, green), bivalent (H3K27me3 and H3K4me1/3, orange) and Polycomb (H3K27me3-only, red) chromatin. (D) Heatmap shows the difference in *HindIII* fragment interaction frequency between cell types as a function of the chromatin state of the interacting regions (rows) and the linear interaction distance (columns, binned distances). Interacting regions that are engaged in long-range promoter interactions, defined as >1Mb, are highlighted by the dashed box. Nearly all (98%) of the long-range interactions were associated with bivalently-marked promoters and these regions have a higher interaction frequency in primed compared to naïve PSCs.

**Figure S6.**
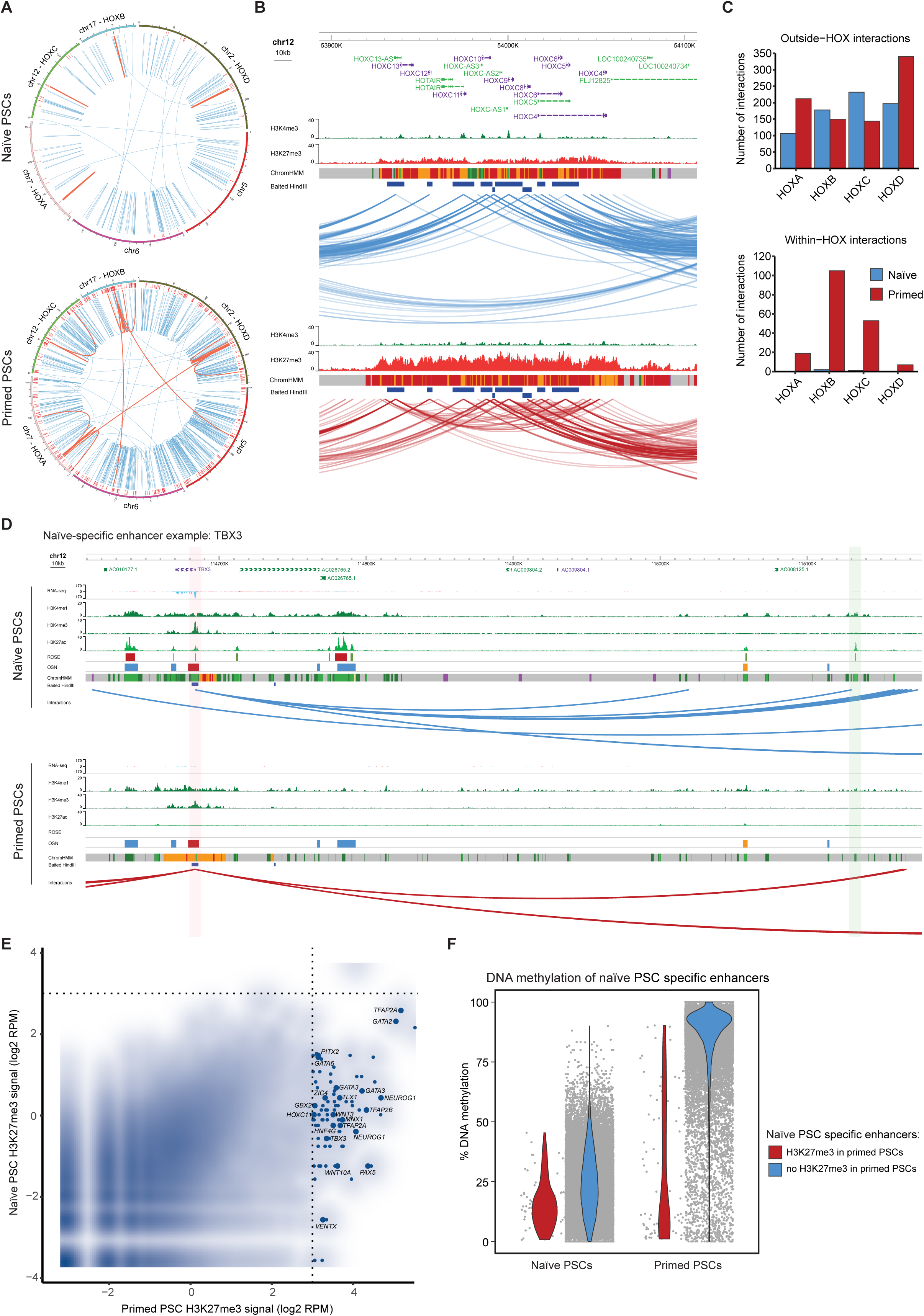
Differences in promoter interactions between human pluripotent states. (A) Circos plots show long-range PCHi-C interactions (blue lines) and interactions with the *HOX* clusters (red lines) for the chromosomes indicated in naïve (upper) and primed (lower) PSCs. The red outer track shows H3K27me3-marked regions. (B) Genome browser representations of the *HOXC* locus in naïve (upper) and primed (lower) PSCs. Tracks shown include all chromatin interactions that have at least one end of an interaction in this region (blue and red lines), baited *HindIII* fragments, and ChIP-Seq data including H3K4me3, H3K27me3 and ChromHMM states. Note the higher number of interactions within the *HOXC* locus in primed compared to naïve PSCs. (C) Charts show the number of ‘outside’ (upper) and ‘within’ (lower) chromatin interactions for the four *HOX* loci in both cell types. ‘Outside’ interactions are when one end of the interaction is within the *HOX* region and the other end of the interaction is outside of the region, and ‘within’ interactions are when both ends of the interaction are within the same *HOX* region. Note the striking difference in the number of ‘within’ interactions, particularly for the *HOXB* and *HOXC* loci, between pluripotent states. (D) Genome browser representation shows an example of a naïve-specific enhancer at the *TBX3* locus. *TBX3* is highly expressed in naïve PSCs, and the *TBX3* promoter (shaded in red box) interacts with a distal active enhancer (shaded in green box) only in naïve PSCs. (E) Smoothed scatter plot shows H3K27me3 levels at naïve-specific enhancer regions in both pluripotent states. Enhancers that gain H3K27me3 in primed PSCs are highlighted and annotated with their nearest gene. (F) Violin plot shows the percent DNA methylation of two classes of naïve-specific enhancers depending on whether those regions gain (red) or do not gain (blue) H3K27me3 after transition to a primed state. Naïve-specific enhancers that acquire H3K27me3 are protected from DNA hypermethylation.

**Figure S7.**
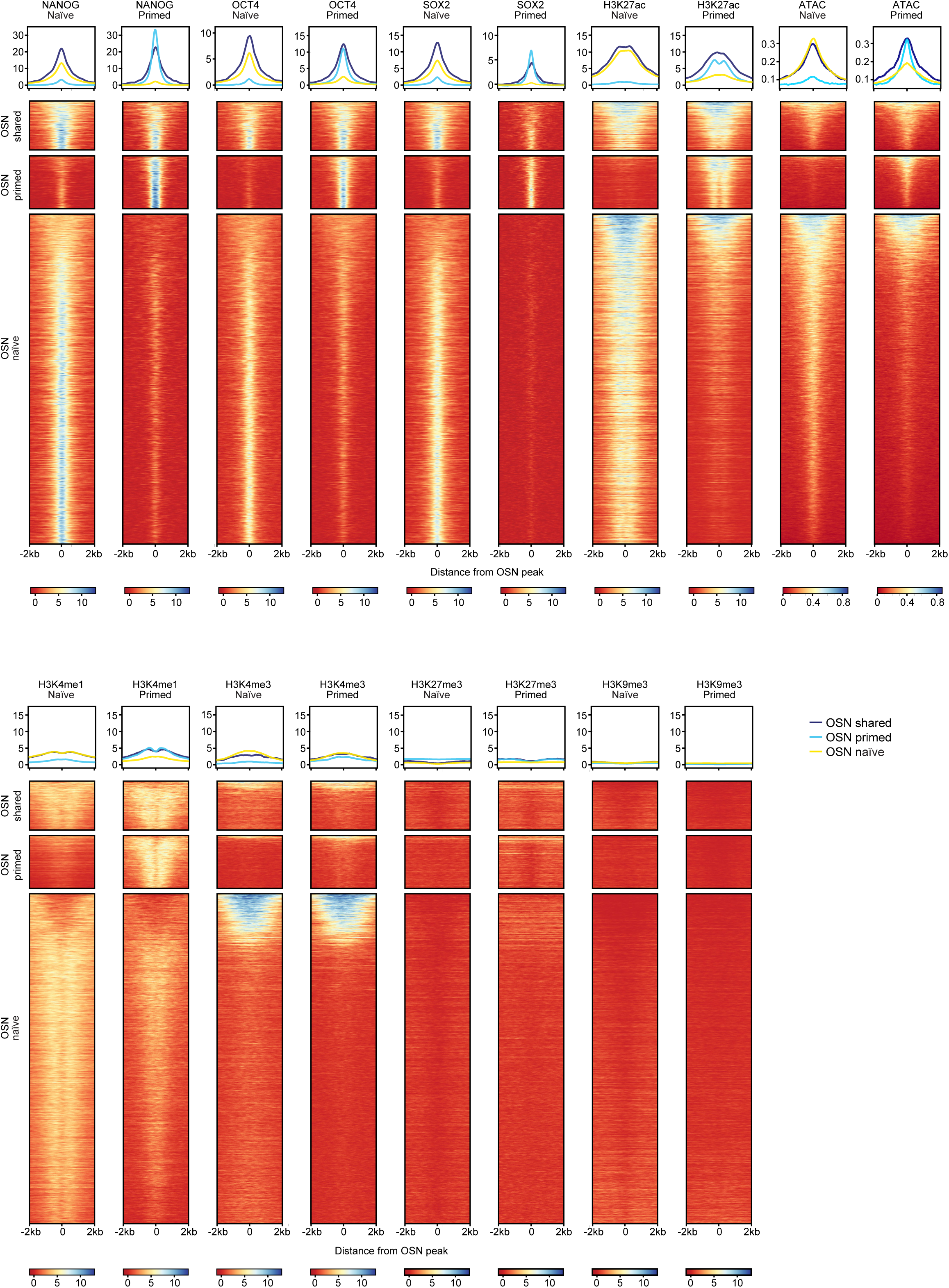
Characterisation of OSN sites in naïve and primed PSCs. ChIP-seq and ATAC-seq data for NANOG, OCT4 and SOX2 were used to categorise OSN sites that are specific to either naïve PSCs (n=13,462) or primed PSCs (n=2,164) or shared between both pluripotent states (n=1,994). Shown are metaplots (top) and heatmaps (bottom) of log2-transformed read counts within a 4 kb window centred on the OSN peak.

